# Guanine content of microRNAs is associated with their tumor suppressive and oncogenic roles in lung and breast cancers

**DOI:** 10.1101/518472

**Authors:** Amit Cohen, Mario Alberto Burgos-Aceves, Yoav Smith

## Abstract

**Background:** microRNAs (miRNAs, miRs) are small noncoding RNAs that negatively regulate gene expression at the post-transcriptional level and fine-tune gene functions. A global repression of miRNAs expression in different types of human tumors, after exposure to cigarette-smoke, or to the hormone estrogen, have been shown to be associated with guanine (G) enrichment in the terminal loops (TLs) of their precursors.

**Methods:** we integrated the G content of miRNA mature forms and precursor miRNA TLs with their described function in the literature, using the PubMed database. Gene Ontology term analysis was used to describe the pathways in which the G-enriched miRNA targets are involved.

**Results:** we show here an association between the relative G enrichment of precursor miRNAs TLs and their tendency to act as tumor suppressor miRs in human lung and breast cancers. Another association was observed between the high G content of the miRNAs 5-mature forms and their tendency to act as oncomiRs.

**Conclusions:** the results support previous findings showing that the G sequence content is an important feature determining miRNA expression and function, and open the way for future cancer investigations in this direction.

## Background

MicroRNAs (miRNAs) are endogenous ~22-nucleotides RNA molecules, that negatively regulate gene expression at the post-transcriptional level and are implicated in the pathogenesis of many human diseases, including cancer, where they can act as tumor suppressive miRNAs (tumor suppressive miRs) or as oncogenic miRNAs (oncomiRs) [1, 2]. Generally, oncomiRs are overexpressed in cancers while tumor suppressive miRs are underexpressed, however, a comprehensive reduction in miRNA was commonly observed in human cancers, where miRNAs showed lower expression levels in tumors and cancer cell lines compared with normal tissues [3–6]. In addition, a widespread repression of miRNA expression has also been reported after exposure to cigarette-smoke (CS) [7–9], treatment with the hormone estrogen [10–12], and c-Myc activation [13]. These aforementioned alterations in miRNA expression can occur as a result of affecting the transcription of miRNA genes [13], miRNA export from the nucleus [14], or at any stage of the miRNA maturation process by modulation of key regulators or components of the miRNA biogenesis pathway, including Drosha and Dicer [15].

Findings suggest that the miRNA terminal loop (TL) is an important platform for different RNA-binding proteins (RBPs) that act as activators or repressors of Drosha and Dicer processing, and selectivity regulate miRNAs by binding to guanine (G)-enriched motifs in the RNA TLs of their precursors [16]. It was shown that miRNAs with the tetra-nucleotide sequence motif GGAG in their TLs were regulated through binding of the RBP Lin28, which interferes with Dicer processing [17], and that the sequence AGGGU in the TL mediates regulation of miRNA biogenesis by the KH-type splicing regulatory protein (KSRP) RBP [18]. It was recently shown that modification of KSRP resulted in the downregulation of a subset of TL-G-rich miRNAs and promoted tumorigenesis [19].

Izzotti and Pulliero showed in their study that the G content of the TLs of miRNAs, which are involved in stress response, is higher than the G content of the other miRNAs [20]. We have recently found, using bioinformatic data analysis of zebrafish, mouse, and human breast cancer cell line, an association between the widespread miRNAs reduction that is observed after estrogen (17β-estradiol; E2) exposure and a high TL-G content in their precursors [21]. In addition, we also showed that similar G enrichment exists in TLs of downregulated miRNAs found in different human cancers [22]. Here, we bioinformatically analyzed the sequences of over 250 human miRNAs, and show the association between miRNAs G-content and their known function as tumor suppressors and oncogenes in lung and breast cancers.

## Materials and Methods

### Literature-data mining

Literature searches were performed in the PubMed literature database for original articles written in the English language focusing on miRNAs and lung or breast cancers. The searches included a specific miRNA term paired with the key words; ‘lung cancer’ or ‘breast cancer’. No restriction was set for the publication date. Only articles showing a role for miRNAs, by using functional studies, were selected. Where applicable, the direct target genes of the investigated miRNAs were retrieved from the above articles.

### Bioinformatic tools

All miRNA precursor sequences were obtained from the Sanger Institute miRBase database version 21 (http://microrna.sanger.ac.uk/sequences/). DAVID v6 functional annotation tool was used to identify enriched GO terms of genes.

### Nucleotide composition analysis

Calculation of nucleotide composition in miRNA precursors was determined using the compseq algorithm (http://emboss.bioinformatics.nl/cgi-bin/emboss/compseq). Input sequences included precursor miRNAs (pre-miRNAs) stem-loops (SLs), TLs, 5- and 3-mature miRNAs of the tumor suppressive miRs and oncomiRs (miRNA lists and their sequences are presented in Supplementary Table 1).

### Statistical data analysis

A one-way ANOVA, post-hoc Tukey HSD Test was performed for comparison between G enrichment in SLs, TLs, and mature miRNAs of the tumor suppressive miRs and oncomiRs. Statistical analysis was performed using the software XLSTAT (Addinsoft Inc., Paris, France).

## Results

Since miRNAs downregulation in cancer is associated with the relative G enrichment of their TL sequences [22], we asked whether there is also a relation between the relative G content of miRNAs and their known function in cancer. For this purpose, 255 human pre-miRNA sequences were retrieved from the miRBase database and used for further analysis. Each sequence was divided into its different structural constituents; the TL and the two mature miRNAs (5-mature and 3-mature forms). The complete list of miRNA sequences is presented in Supplementary Table 1. For each of these miRNAs the complete pre-miRNA, the TL, and the 5 and 3-mature miRNA sequences were analyzed for evaluation of nucleotide composition (Supplementary Table 1). Next, we filtered the miRNA list and selected those with relatively high G content (more than 35%) or low G content (less than 15%) in their TL and 5-mature sequences, and for the resulted 105 miRNAs, we searched for known functions in lung and breast cancers, by mining publicly available data in the PubMed database. The search resulted in 420 articles; 109 studied on oncomiRs and 311 on tumor suppressive miRs.

The results show that when presenting the number of the articles respectively to the G percentage of miRNAs TLs, tumor suppressive miRs are found to be more G-enriched in their TLs, while oncomiRs show the opposite trend (Figure 1A). When presenting the number of the articles respectively to the G percentage of 5-mature miRNAs, the oncomiRs appear to be more G-enriched in their 5-mature forms (Figure 1B). From this list, we selected only those miRNAs that showed a tendency to act as either tumor suppressive miRs or oncomiRs, in both lung and breast cancers. A list of 84 miRNAs was obtained and sub-grouped into oncomiRs (25 miRNAs) and tumor suppressive miRs (59 miRNAs) (Supplementary Table 2).

**Figure 1.**
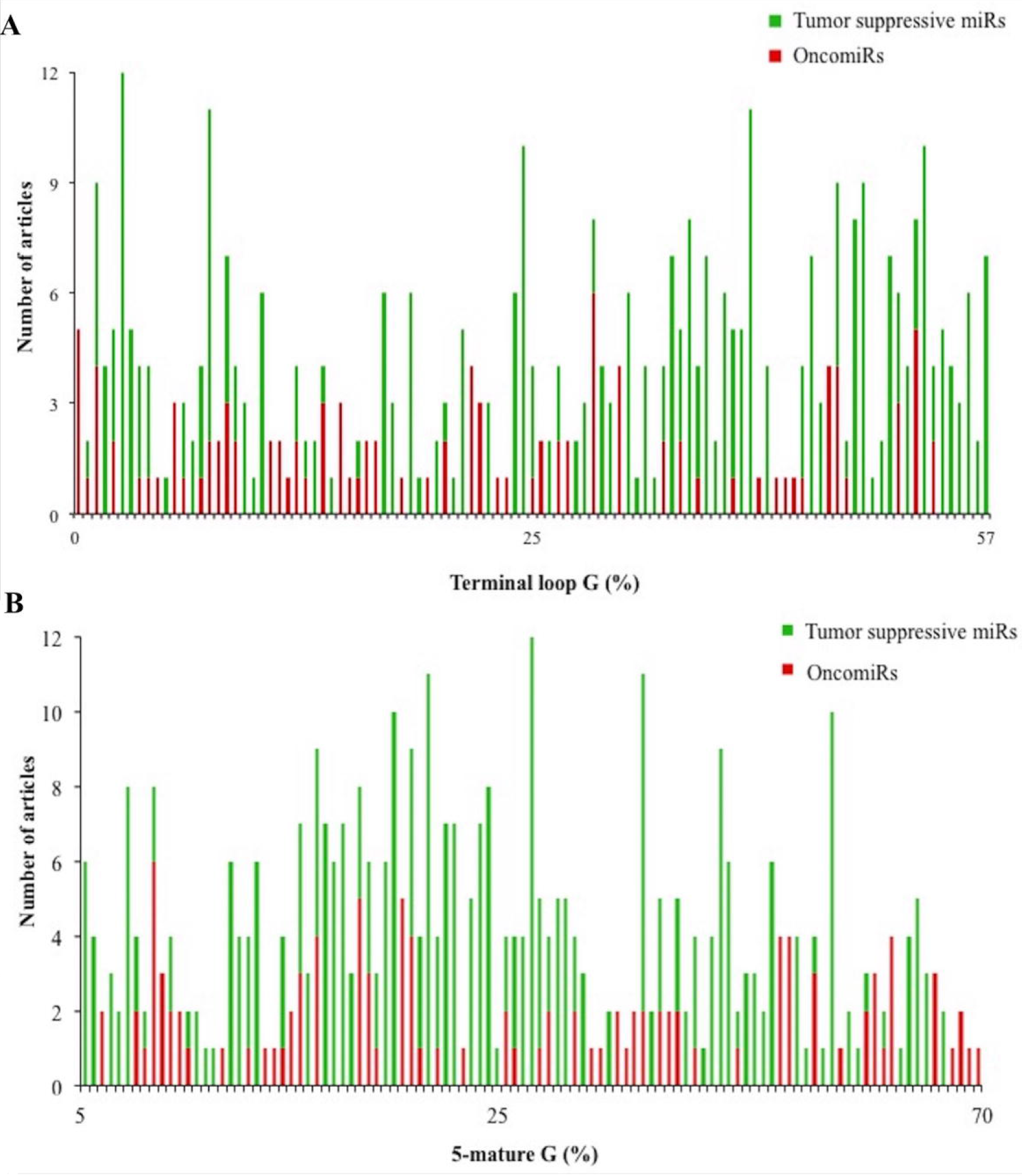
Number of PubMed articles describing the function of tumor suppressive miRs and oncomiRs relative to (a). TL G enrichment. (b). 5-mature G enrichment.

The results show that the TLs of the tumor suppressive miRs group are more enriched in G (1.33 fold) than the group of oncomiRs (Figure 2). No such enrichment was observed when the same analysis was conducted with the complete pre-miRNA SL sequences, or with the 5 and 3-mature miRNAs (Figure 2). Moreover, this enrichment is even more prominent when analyzing the dual G (GG) content, where the TLs of the tumor suppressive miRs group are more enriched in GG (2.1 fold) than the group of oncomiRs (Figure 2). Also here, no such enrichment was observed when the same analysis was conducted with the complete SLs, despite this time the GG content of the 3-mature miRNAs was also slightly enriched (1.32 fold). Also in the case of triple G (GGG), only the TLs of the tumor suppressive miRs group are more enriched in G (1.34 fold) than the group of oncomiRs (Figure 2). Interestingly, the G content of the 5-mature miRNAs has shown the opposite trend than the TLs, and was slightly enriched in oncomiRs relative to tumor suppressive miRs (0.87 fold) (Figure 2), and this trend of enrichment in the 5-mature forms was even more pronounced when looking at the GG (0.65 fold) and GGG (0.39 fold) content (Figure 2). Statistical analysis revealed a significant difference between TLs and 5-mature miRNAs G, GG and GGG content of the tumor suppressive miRs relative to oncomiRs (one-way ANOVA, *P*=0.01). Together, the above results show an association between high TLs and 5-mature miRNAs G content and their tendency to act either as tumor suppressive miRs or oncomiRs, respectively.

**Figure 2.**
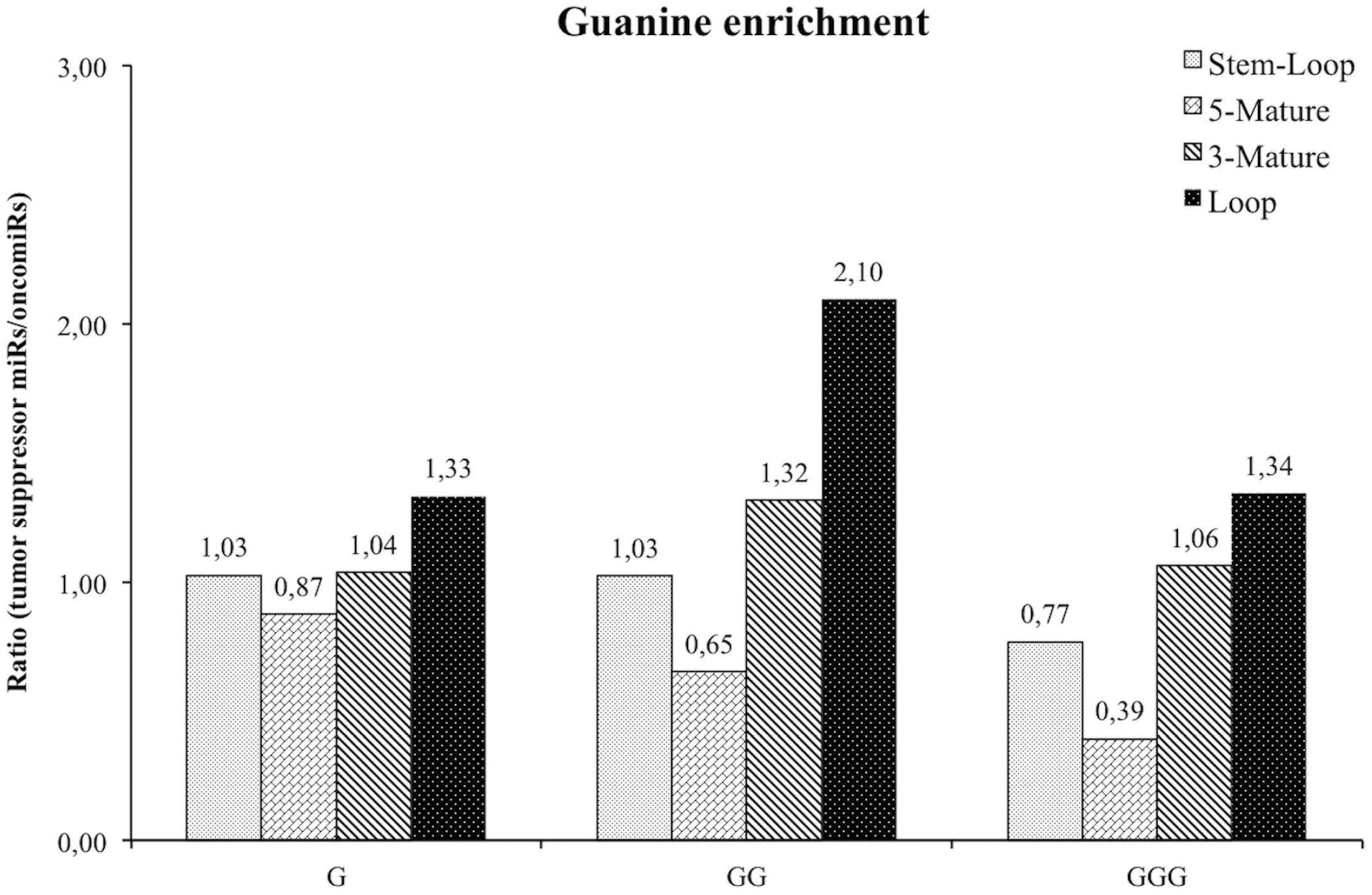
G enrichment of tumor suppressive miRs relative to oncomiRs. Single, dual and triple G were calculated in SLs, TLs, 5 and 3-mature miRNAs involved in lung and breast cancers.

In order to define the function of those tumor suppressive miRs, which have relatively high G content in their TLs, we selected the 30 most TL-G-enriched miRNAs, (over 35% G), retrieved their identified direct target genes from the PubMed articles (Supplementary Table 2), and searched for biological processes enrichment. Functional annotation was performed using the DAVID tool, and among Gene Ontology (GO) biological processes; negative regulation of cell death, regulation of signal transduction, and positive regulation of cell migration, were the most significant ones in lung and breast cancers (Figure 3). The results show that a large number of target genes of these tumor suppressive miRs are known proto-oncogenes (MYC, MYCN, EGFR, ERBB2, ERBB3, ERBB4, AKT1, CCND1, KRAS, SRC, BCL2, PIK3CA, MET, AXL, PDGFRB, JAK2, HMGA2, MYB, AGR2, PRKCA, PIM1, RAF1, RHOA, MKL1, SALL4) (Supplementary Table 3). In several cases, these oncogenes were common targets to multiple miRNAs, such as in KRAS (miR-30c, −143, −193a, −200c, let-7a), CCND1 (miR-34b, −145a, −195, let-7e), and EGFR (miR-34a, −143, −218) (Supplementary Table 3).

**Figure 3.**
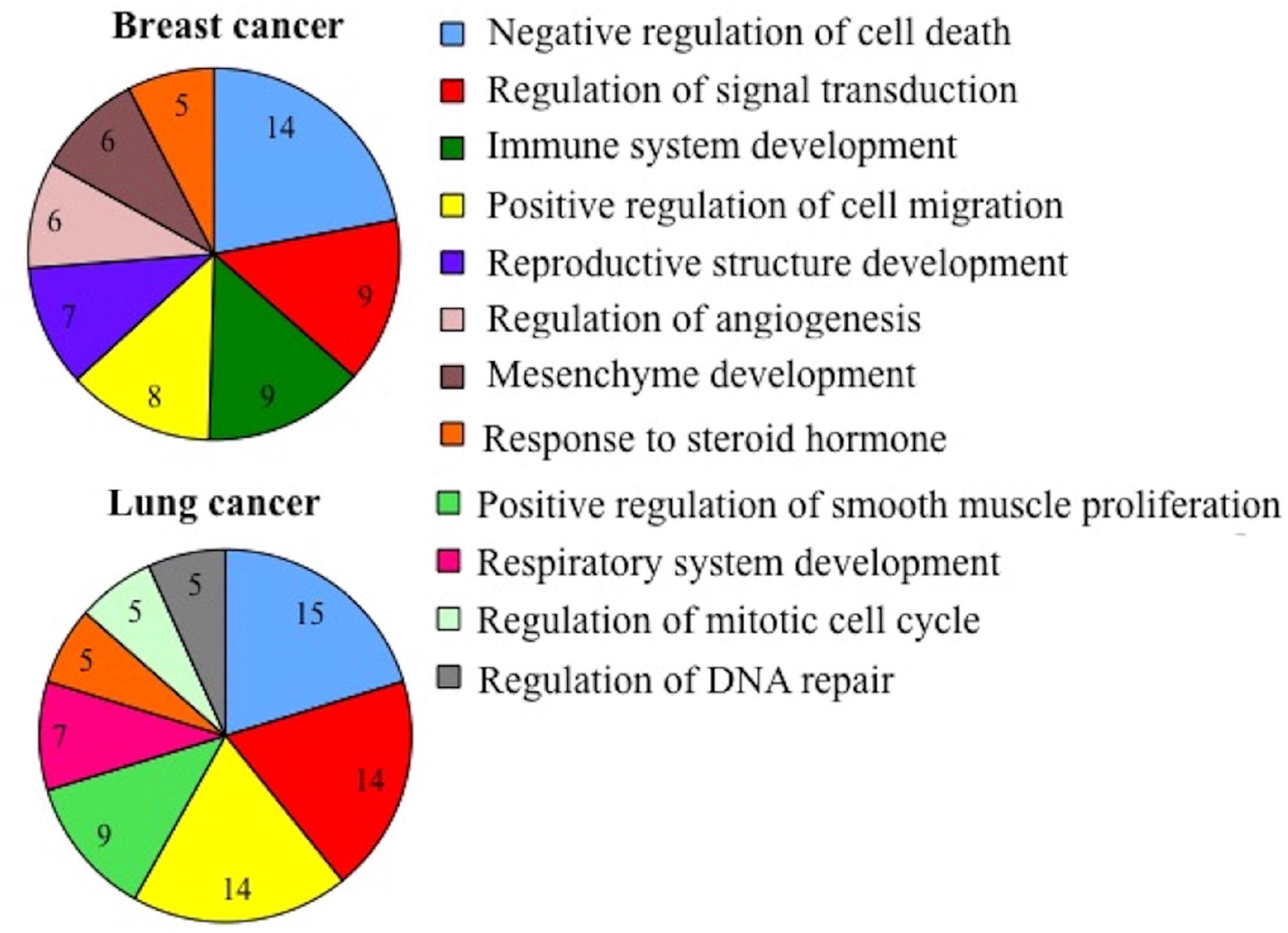
Enriched GO terms of the target genes of G-enriched tumor suppressive miRs. Shown are significant −log2 Benjamini p-values of the biological process terms (level 5) of the 30 most TL-G-enriched tumor suppressive miRs target genes, as were identified using the DAVID functional annotation tool.

## Discussion

Over the past two decades, it has become increasingly apparent that loss- or gain-of-function of specific miRNAs contributes to cellular transformation and tumorigenesis [23]. OncomiRs typically function to promote cell growth, inhibit apoptosis and control the cell cycle, while tumor suppressive miRs function to inhibit cell growth and induce apoptosis [24], and there are several examples in which a specific miRNA can act either as a tumor suppressor or an oncomiR, depending on cellular context and type of cancer [25]. However, a global miRNA downregulation in cancer was also observed, and therefore, understanding the mechanisms driving miRNA downregulation is important in uncovering the regulatory role of miRNAs in cancer biology [26].

The results presented here support previous studies showing the potential importance of TL-G content to the regulation of miRNAs expression and function [19–22]. We have previously suggested that estrogens metabolites inside the lungs, as a result of CS exposure, can potentially cause the observed widespread downregulation of miRNA expression, and contribute to lung tumor development [27, 28]. Estrogen metabolism generates highly reactive metabolites, mostly catechol estrogen-3,4-quinones, which form carcinogen depurinating G adducts [29]. Moreover, oxidative metabolites of estrogens can cause oxidative damage to G, because of its lowest oxidation potential, by forming 8-oxo-dG (8-Oxo-2′-deoxyguanosine), which eventually leads to carcinogenesis [30]. Repressed miRNAs in lung and breast cancers were shown to be the most enriched in TLs G [22], and the relation of estrogen to the development of these types of cancer is well documented [27, 30, 31]. Furthermore, like in the current study, also in the aforementioned studies, dual GG enrichment in miRNA TLs was even more pronounced [20–22]. Remarkably, experimental studies have shown that sequences with repeated G bases (GG or GGG) show higher reactivity toward oxidation than isolated G bases [32].

Despite analyzing during this study relatively small fraction (~14%) of the total number of currently identified human miRNAs, our results indicate that the number of tumor suppressive miRs is greater than oncomiRs. This is in agreement with the finding that many of the known mutated genes are oncogenes [33], and with the general consideration of miRNAs as safeguards of the genome [24]. Of note, a remarkable redundancy was observed between the target oncogenes of the tumor suppressive miRs (e.g. KRAS). Synergetic effects of functionally related tumor suppressive miRs, that share common targets and control similar processes, were shown before [34].

As indicated above, tumor suppressive miRs tended to be more G-enriched in their TLs. G enrichment in tumor suppressive miRs TLs could affect binding of RBPs such as Lin28, KSRP, and the heteronuclear ribonucleoprotein A1 (hnRNP A1), which compete with KSRP for their common G-rich target sequence [35]. Notably, hnRNPA2B1, another splicing factor, binds exosomal miRNAs through the recognition of G-enriched motifs to control their loading into exosomes [36]; small membranous vesicles which enable genetic exchange between cells, can transfer functional miRNAs to recipient cells and consequently downregulate the expression of their target genes [37]. It is noteworthy that sumoylation modification controls the binding of both KSRP and hnRNPA2B1 to miRNAs [19, 36].

OncomiRs, on the contrary, have lower G content in their TLs, and are relatively more G-enriched in their 5-mature miRNA forms. The differences in G enrichment shown here between the 5 and 3-mature miRNAs might be attributed to the differences between the functional guide strand and the passenger strand of mature miRNAs, as one of the characteristics of human miRNA guide strands is excess of purines [38]. This observation could also be related to the process of sorting miRNAs into the exosomes, as it was shown that G-rich sequence is a dominant feature of exosome-dominant miRNAs, suggesting the possibility that RBP-mediated translocation of cellular miRNAs into exosome cargos occurs by G-recognition [39]. Indeed, oncogenic exosomal miRNAs (miR-17, −21, −106a, −155, −191) were highly induced in lung cancer [40, 41], and chemoresistance of breast cancer cells was recently shown to occur through the G-enriched miR-155 oncomiR exosomes delivery [42].

## Conclusions

Taken together, the current and previously published results suggest that G content of miRNAs is an important feature determining their expression and function. MiRNA TL-G enrichment may have important role in the carcinogenic process by affecting the observed global downregulation of tumor suppressive miRs, which cause induction of their target oncogenes, and commit cells towards carcinogenesis. On the other hand, high G content in the oncomiRs 5-mature sequences may function in guide-strand selection and miRNA targeting into the exosomes. Elucidating the molecular mechanisms that involve miRNAs G can have major implications for cancer research, therapy and prevention [43, 44].

## Acknowledgements

None

## Supplementary Material

**Supplementary Table 1:** SLs, TLs, 5 and 3-mature miRNA sequences, and their G enrichment. Shown are 255 miRNA sequences retrieved from the miRBase database. G enrichment of sequences represents the ratio of G number relative to total nucleotide number.

**Supplementary Table 2:** Tumor suppressive miRs, oncomiRs, and the references from PubMed Database describing their function in lung and breast cancers. When search in PubMed had no results it appears as unknown function.

**Supplementary Table 3:** G-enriched tumor suppressive miRs target genes. Shown are 30 most TL-G-enriched tumor suppressive miRs and their direct target genes in lung and breast cancers, as were described in the references from PubMed Database (Supplementary Table 2).

Contributions

1. Conception and design: AC
2. Administrative support: MAB-A
3. Provision of study materials or patients: AC, MAB-A
4. Collection and assembly of data: All authors
5. Data analysis and interpretation: AC, YS
6. Manuscript writing: All authors
7. Final approval of manuscript: All authors.

