## Supplementary material for "Guanine content of microRNAs is associated with their tumor suppressive and oncogenic roles in lung and breast cancers": Supp Table 1

Supplementary Table 1. SLs, TLs, 5 and 3-mature miRNA sequences, and their G enrichment. Shown are 255 miRNA sequences retrieved from the miRBase database. G enrichment (%) of sequences represents the ratio of G number relative to total nucleotide number.

| **miRNA** | **Stem-loop** | **G enrichment** | **Loop** | **G enrichment** | **5-mature** | **G enrichment** | **3-mature** | **G enrichment** |
| --- | --- | --- | --- | --- | --- | --- | --- | --- |
| hsa-let-7a-1 | UGGGAUGAGGUAGUAGGUUGUAUAGUUUUAGGGUCACACCCACCACUGGGAGAUAACUAUACAAUCUACUGUCUUUCCUA | 24 | UUAGGGUCACACCCACCACUGGGAGAUAA | 24 | UGAGGUAGUAGGUUGUAUAGUU | 36 | CUAUACAAUCUACUGUCUUUC | 5 |
| hsa-let-7a-2 | AGGUUGAGGUAGUAGGUUGUAUAGUUUAGAAUUACAUCAAGGGAGAUAACUGUACAGCCUCCUAGCUUUCCU | 25 | UAGAAUUACAUCAAGGGAGAUAA | 22 | UGAGGUAGUAGGUUGUAUAGUU | 36 | CUGUACAGCCUCCUAGCUUUCC | 14 |
| hsa-let-7a-3 | GGGUGAGGUAGUAGGUUGUAUAGUUUGGGGCUCUGCCCUGCUAUGGGAUAACUAUACAAUCUACUGUCUUUCCU | 28 | UGGGGCUCUGCCCUGCUAUGGGAUAA | 35 | UGAGGUAGUAGGUUGUAUAGUU | 36 | CUAUACAAUCUACUGUCUUUC | 5 |
| hsa-let-7b | CGGGGUGAGGUAGUAGGUUGUGUGGUUUCAGGGCAGUGAUGUUGCCCCUCGGAAGAUAACUAUACAACCUACUGCCUUCCCUG | 31 | UCAGGGCAGUGAUGUUGCCCCUCGGAAGAUAA | 31 | UGAGGUAGUAGGUUGUGUGGUU | 45 | CUAUACAACCUACUGCCUUCCC | 5 |
| hsa-let-7c | GCAUCCGGGUUGAGGUAGUAGGUUGUAUGGUUUAGAGUUACACCCUGGGAGUUAACUGUACAACCUUCUAGCUUUCCUUGGAGC | 29 | UAGAGUUACACCCUGGGAGUUAA | 26 | UGAGGUAGUAGGUUGUAUGGUU | 41 | CUGUACAACCUUCUAGCUUUCC | 9 |
| hsa-let-7d | CCUAGGAAGAGGUAGUAGGUUGCAUAGUUUUAGGGCAGGGAUUUUGCCCACAAGGAGGUAACUAUACGACCUGCUGCCUUUCUUAGG | 30 | UUAGGGCAGGGAUUUUGCCCACAAGGAGGUAA | 34 | AGAGGUAGUAGGUUGCAUAGUU | 36 | CUAUACGACCUGCUGCCUUUCU | 14 |
| hsa-let-7e | CCCGGGCUGAGGUAGGAGGUUGUAUAGUUGAGGAGGACACCCAAGGAGAUCACUAUACGGCCUCCUAGCUUUCCCCAGG | 32 | GAGGAGGACACCCAAGGAGAUCA | 35 | UGAGGUAGGAGGUUGUAUAGUU | 41 | CUAUACGGCCUCCUAGCUUUCC | 14 |
| hsa-let-7f-1 | UCAGAGUGAGGUAGUAGAUUGUAUAGUUGUGGGGUAGUGAUUUUACCCUGUUCAGGAGAUAACUAUACAAUCUAUUGCCUUCCCUGA | 25 | GUGGGGUAGUGAUUUUACCCUGUUCAGGAGAUAA | 32 | UGAGGUAGUAGAUUGUAUAGUU | 32 | CUAUACAAUCUAUUGCCUUCCC | 5 |
| hsa-let-7f-2 | UGUGGGAUGAGGUAGUAGAUUGUAUAGUUUUAGGGUCAUACCCCAUCUUGGAGAUAACUAUACAGUCUACUGUCUUUCCCACG | 24 | UUAGGGUCAUACCCCAUCUUGGAGAUAA | 21 | UGAGGUAGUAGAUUGUAUAGUU | 32 | CUAUACAGUCUACUGUCUUUCC | 9 |
| hsa-let-7g | AGGCUGAGGUAGUAGUUUGUACAGUUUGAGGGUCUAUGAUACCACCCGGUACAGGAGAUAACUGUACAGGCCACUGCCUUGCCA | 29 | UGAGGGUCUAUGAUACCACCCGGUACAGGAGAUAA | 29 | UGAGGUAGUAGUUUGUACAGUU | 32 | CUGUACAGGCCACUGCCUUGC | 24 |
| hsa-let-7i | CUGGCUGAGGUAGUAGUUUGUGCUGUUGGUCGGGUUGUGACAUUGCCCGCUGUGGAGAUAACUGCGCAAGCUACUGCCUUGCUA | 33 | GGUCGGGUUGUGACAUUGCCCGCUGUGGAGAUAA | 38 | UGAGGUAGUAGUUUGUGCUGUU | 36 | CUGCGCAAGCUACUGCCUUGCU | 23 |
| hsa-mir-100 | CCUGUUGCCACAAACCCGUAGAUCCGAACUUGUGGUAUUAGUCCGCACAAGCUUGUAUCUAUAGGUAUGUGUCUGUUAGG | 24 | GUAUUAGUCCGCA | 23 | AACCCGUAGAUCCGAACUUGUG | 23 | CAAGCUUGUAUCUAUAGGUAUG | 23 |
| hsa-mir-101-1 | UGCCCUGGCUCAGUUAUCACAGUGCUGAUGCUGUCUAUUCUAAAGGUACAGUACUGUGAUAACUGAAGGAUGGCA | 25 | GUCUAUUCUAAAGG | 21 | CAGUUAUCACAGUGCUGAUGCU | 23 | UACAGUACUGUGAUAACUGAA | 19 |
| hsa-mir-103a-2 | UUGUGCUUUCAGCUUCUUUACAGUGCUGCCUUGUAGCAUUCAGGUCAAGCAGCAUUGUACAGGGCUAUGAAAGAACCA | 23 | UAGCAUUCAGGUC | 23 | AGCUUCUUUACAGUGCUGCCUUG | 22 | AGCAGCAUUGUACAGGGCUAUGA | 30 |
| hsa-mir-105-1 | UGUGCAUCGUGGUCAAAUGCUCAGACUCCUGUGGUGGCUGCUCAUGCACCACGGAUGUUUGAGCAUGUGCUACGGUGUCUA | 30 | GGCUGCUCAUGCACC | 27 | UCAAAUGCUCAGACUCCUGUGGU | 22 | ACGGAUGUUUGAGCAUGUGCUA | 32 |
| hsa-mir-105-2 | UGUGCAUCGUGGUCAAAUGCUCAGACUCCUGUGGUGGCUGCUUAUGCACCACGGAUGUUUGAGCAUGUGCUAUGGUGUCUA | 30 | GGCUGCUUAUGCACC | 27 | UCAAAUGCUCAGACUCCUGUGGU | 22 | ACGGAUGUUUGAGCAUGUGCUA | 32 |
| hsa-mir-106a | CCUUGGCCAUGUAAAAGUGCUUACAGUGCAGGUAGCUUUUUGAGAUCUACUGCAAUGUAAGCACUUCUUACAUUACCAUGG | 21 | CUUUUUGAGAUCUA | 14 | AAAAGUGCUUACAGUGCAGGUAG | 30 | CUGCAAUGUAAGCACUUCUUAC | 14 |
| hsa-mir-106b | CCUGCCGGGGCUAAAGUGCUGACAGUGCAGAUAGUGGUCCUCUCCGUGCUACCGCACUGUGGGUACUUGCUGCUCCAGCAGG | 32 | AGUGGUCCUCUCCGUGCUA | 26 | UAAAGUGCUGACAGUGCAGAU | 29 | CCGCACUGUGGGUACUUGCUGC | 32 |
| hsa-mir-10a | GAUCUGUCUGUCUUCUGUAUAUACCCUGUAGAUCCGAAUUUGUGUAAGGAAUUUUGUGGUCACAAAUUCGUAUCUAGGGGAAUAUGUAGUUGACAUAAACACUCCGCUCU | 21 | UAAGGAAUUUUGUGGUCA | 28 | UACCCUGUAGAUCCGAAUUUGUG | 22 | CAAAUUCGUAUCUAGGGGAAUA | 23 |
| hsa-mir-10b | CCAGAGGUUGUAACGUUGUCUAUAUAUACCCUGUAGAACCGAAUUUGUGUGGUAUCCGUAUAGUCACAGAUUCGAUUCUAGGGGAAUAUAUGGUCGAUGCAAAAACUUCA | 23 | UGGUAUCCGUAUAGUC | 25 | UACCCUGUAGAACCGAAUUUGUG | 22 | ACAGAUUCGAUUCUAGGGGAAU | 27 |
| hsa-mir-1-1 | UGGGAAACAUACUUCUUUAUAUGCCCAUAUGGACCUGCUAAGCUAUGGAAUGUAAAGAAGUAUGUAUCUCA | 20 | AUGGACCUGCUAAGCUA | 24 | ACAUACUUCUUUAUAUGCCCAU | 5 | UGGAAUGUAAAGAAGUAUGUAU | 27 |
| hsa-mir-122 | CCUUAGCAGAGCUGUGGAGUGUGACAAUGGUGUUUGUGUCUAAACUAUCAAACGCCAUUAUCACACUAAAUAGCUACUGCUAGGC | 22 | UGUCUAAACUAUCA | 7 | UGGAGUGUGACAAUGGUGUUUG | 41 | AACGCCAUUAUCACACUAAAUA | 5 |
| hsa-mir-124-1 | AGGCCUCUCUCUCCGUGUUCACAGCGGACCUUGAUUUAAAUGUCCAUACAAUUAAGGCACGCGGUGAAUGCCAAGAAUGGGGCUG | 26 | UUAAAUGUCCAUACAAU | 6 | CGUGUUCACAGCGGACCUUGAU | 27 | UAAGGCACGCGGUGAAUGCC | 35 |
| hsa-mir-124-2 | AUCAAGAUUAGAGGCUCUGCUCUCCGUGUUCACAGCGGACCUUGAUUUAAUGUCAUACAAUUAAGGCACGCGGUGAAUGCCAAGAGCGGAGCCUACGGCUGCACUUGAA | 26 | UUAAUGUCAUACAAU | 7 | CGUGUUCACAGCGGACCUUGAU | 27 | UAAGGCACGCGGUGAAUGCC | 35 |
| hsa-mir-124-3 | UGAGGGCCCCUCUGCGUGUUCACAGCGGACCUUGAUUUAAUGUCUAUACAAUUAAGGCACGCGGUGAAUGCCAAGAGAGGCGCCUCC | 28 | UUAAUGUCUAUACAAU | 6 | CGUGUUCACAGCGGACCUUGAU | 27 | UAAGGCACGCGGUGAAUGCC | 35 |
| hsa-mir-125a | UGCCAGUCUCUAGGUCCCUGAGACCCUUUAACCUGUGAGGACAUCCAGGGUCACAGGUGAGGUUCUUGGGAGCCUGGCGUCUGGCC | 31 | GGACAUCCAGGGUC | 36 | UCCCUGAGACCCUUUAACCUGUGA | 17 | ACAGGUGAGGUUCUUGGGAGCC | 41 |
| hsa-mir-125b-1 | UGCGCUCCUCUCAGUCCCUGAGACCCUAACUUGUGAUGUUUACCGUUUAAAUCCACGGGUUAGGCUCUUGGGAGCUGCGAGUCGUGCU | 26 | UGUUUACCGUUUAAAUCC | 11 | UCCCUGAGACCCUAACUUGUGA | 18 | ACGGGUUAGGCUCUUGGGAGCU | 41 |
| hsa-mir-125b-2 | ACCAGACUUUUCCUAGUCCCUGAGACCCUAACUUGUGAGGUAUUUUAGUAACAUCACAAGUCAGGCUCUUGGGACCUAGGCGGAGGGGA | 26 | GGUAUUUUAGUAACA | 20 | UCCCUGAGACCCUAACUUGUGA | 18 | UCACAAGUCAGGCUCUUGGGAC | 27 |
| hsa-mir-126 | CGCUGGCGACGGGACAUUAUUACUUUUGGUACGCGCUGUGACACUUCAAACUCGUACCGUGAGUAAUAAUGCGCCGUCCACGGCA | 26 | CUGUGACACUUCAAAC | 13 | CAUUAUUACUUUUGGUACGCG | 19 | UCGUACCGUGAGUAAUAAUGCG | 27 |
| hsa-mir-127 | UGUGAUCACUGUCUCCAGCCUGCUGAAGCUCAGAGGGCUCUGAUUCAGAAAGAUCAUCGGAUCCGUCUGAGCUUGGCUGGUCGGAAGUCUCAUCAUC | 27 | UCAGAAAGAUCA | 17 | CUGAAGCUCAGAGGGCUCUGAU | 32 | UCGGAUCCGUCUGAGCUUGGCU | 32 |
| hsa-mir-128-1 | UGAGCUGUUGGAUUCGGGGCCGUAGCACUGUCUGAGAGGUUUACAUUUCUCACAGUGAACCGGUCUCUUUUUCAGCUGCUUC | 27 | GGUUUACAUUUC | 17 | CGGGGCCGUAGCACUGUCUGAGA | 39 | UCACAGUGAACCGGUCUCUUU | 19 |
| hsa-mir-128-2 | UGUGCAGUGGGAAGGGGGGCCGAUACACUGUACGAGAGUGAGUAGCAGGUCUCACAGUGAACCGGUCUCUUUCCCUACUGUGUC | 33 | GUGAGUAGCAGGUC | 43 | GGGGGCCGAUACACUGUACGAGA | 39 | UCACAGUGAACCGGUCUCUUU | 19 |
| hsa-mir-129-1 | GGAUCUUUUUGCGGUCUGGGCUUGCUGUUCCUCUCAACAGUAGUCAGGAAGCCCUUACCCCAAAAAGUAUCU | 22 | UGUUCCUCUCAACAGUAGUCAGG | 22 | CUUUUUGCGGUCUGGGCUUGC | 33 | AAGCCCUUACCCCAAAAAGUAU | 9 |
| hsa-mir-129-2 | UGCCCUUCGCGAAUCUUUUUGCGGUCUGGGCUUGCUGUACAUAACUCAAUAGCCGGAAGCCCUUACCCCAAAAAGCAUUUGCGGAGGGCG | 26 | UGUACAUAACUCAAUAGCCGG | 19 | CUUUUUGCGGUCUGGGCUUGC | 33 | AAGCCCUUACCCCAAAAAGCAU | 9 |
| hsa-mir-130a | UGCUGCUGGCCAGAGCUCUUUUCACAUUGUGCUACUGUCUGCACCUGUCACUAGCAGUGCAAUGUUAAAAGGGCAUUGGCCGUGUAGUG | 27 | ACCUGUCACUAG | 17 | UUCACAUUGUGCUACUGUCUGC | 18 | CAGUGCAAUGUUAAAAGGGCAU | 27 |
| hsa-mir-130b | GGCCUGCCCGACACUCUUUCCCUGUUGCACUACUAUAGGCCGCUGGGAAGCAGUGCAAUGAUGAAAGGGCAUCGGUCAGGUC | 29 | UAUAGGCCGCUGGGAAG | 41 | ACUCUUUCCCUGUUGCACUAC | 10 | CAGUGCAAUGAUGAAAGGGCAU | 32 |
| hsa-mir-132 | CCGCCCCCGCGUCUCCAGGGCAACCGUGGCUUUCGAUUGUUACUGUGGGAACUGGAGGUAACAGUCUACAGCCAUGGUCGCCCCGCAGCACGCCCACGCGC | 29 | GUGGGAACUGGAGG | 57 | ACCGUGGCUUUCGAUUGUUACU | 23 | UAACAGUCUACAGCCAUGGUCG | 23 |
| hsa-mir-133a-1 | ACAAUGCUUUGCUAGAGCUGGUAAAAUGGAACCAAAUCGCCUCUUCAAUGGAUUUGGUCCCCUUCAACCAGCUGUAGCUAUGCAUUGA | 20 | CGCCUCUUCAAUGGA | 20 | AGCUGGUAAAAUGGAACCAAAU | 23 | UUUGGUCCCCUUCAACCAGCUG | 18 |
| hsa-mir-133a-2 | GGGAGCCAAAUGCUUUGCUAGAGCUGGUAAAAUGGAACCAAAUCGACUGUCCAAUGGAUUUGGUCCCCUUCAACCAGCUGUAGCUGUGCAUUGAUGGCGCCG | 27 | CGACUGUCCAAUGGA | 27 | AGCUGGUAAAAUGGAACCAAAU | 23 | UUUGGUCCCCUUCAACCAGCUG | 18 |
| hsa-mir-134 | CAGGGUGUGUGACUGGUUGACCAGAGGGGCAUGCACUGUGUUCACCCUGUGGGCCACCUAGUCACCAACCCUC | 30 | CAUGCACUGUGUUCAC | 19 | UGUGACUGGUUGACCAGAGGGG | 45 | CCUGUGGGCCACCUAGUCACCAA | 22 |
| hsa-mir-135a-1 | AGGCCUCGCUGUUCUCUAUGGCUUUUUAUUCCUAUGUGAUUCUACUGCUCACUCAUAUAGGGAUUGGAGCCGUGGCGCACGGCGGGGACA | 28 | UUCUACUGCUCACUCA | 6 | UAUGGCUUUUUAUUCCUAUGUGA | 17 | UAUAGGGAUUGGAGCCGUGGCG | 45 |
| hsa-mir-135b | CACUCUGCUGUGGCCUAUGGCUUUUCAUUCCUAUGUGAUUGCUGUCCCAAACUCAUGUAGGGCUAAAAGCCAUGGGCUACAGUGAGGGGCGAGCUCC | 27 | UUGCUGUCCCAAACUC | 13 | UAUGGCUUUUCAUUCCUAUGUGA | 17 | AUGUAGGGCUAAAAGCCAUGGG | 36 |
| hsa-mir-136 | UGAGCCCUCGGAGGACUCCAUUUGUUUUGAUGAUGGAUUCUUAUGCUCCAUCAUCGUCUCAAAUGAGUCUUCAGAGGGUUCU | 23 | UUCUUAUGCUC | 9 | ACUCCAUUUGUUUUGAUGAUGGA | 22 | CAUCAUCGUCUCAAAUGAGUCU | 14 |
| hsa-mir-138-1 | CCCUGGCAUGGUGUGGUGGGGCAGCUGGUGUUGUGAAUCAGGCCGUUGCCAAUCAGAGAACGGCUACUUCACAACACCAGGGCCACACCACACUACAGG | 30 | UUGCCAAUCAGAGAACG | 24 | AGCUGGUGUUGUGAAUCAGGCCG | 39 | GCUACUUCACAACACCAGGGCC | 18 |
| hsa-mir-138-2 | CGUUGCUGCAGCUGGUGUUGUGAAUCAGGCCGACGAGCAGCGCAUCCUCUUACCCGGCUAUUUCACGACACCAGGGUUGCAUCA | 27 | ACGAGCAGCGCAUCCUCUUACCCG | 21 | AGCUGGUGUUGUGAAUCAGGCCG | 39 | GCUAUUUCACGACACCAGGGUU | 23 |
| hsa-mir-139 | GUGUAUUCUACAGUGCACGUGUCUCCAGUGUGGCUCGGAGGCUGGAGACGCGGCCCUGUUGGAGUAAC | 35 | GUGGCUCGGAGGC | 54 | UCUACAGUGCACGUGUCUCCAGU | 22 | UGGAGACGCGGCCCUGUUGGAGU | 43 |
| hsa-mir-140 | UGUGUCUCUCUCUGUGUCCUGCCAGUGGUUUUACCCUAUGGUAGGUUACGUCAUGCUGUUCUACCACAGGGUAGAACCACGGACAGGAUACCGGGGCACC | 27 | GUUACGUCAUGCUGUUC | 24 | CAGUGGUUUUACCCUAUGGUAG | 27 | UACCACAGGGUAGAACCACGG | 29 |
| hsa-mir-141 | CGGCCGGCCCUGGGUCCAUCUUCCAGUACAGUGUUGGAUGGUCUAAUUGUGAAGCUCCUAACACUGUCUGGUAAAGAUGGCUCCCGGGUGGGUUC | 31 | UGGUCUAAUUGUGAAGCUCC | 25 | CAUCUUCCAGUACAGUGUUGGA | 23 | UAACACUGUCUGGUAAAGAUGG | 27 |
| hsa-mir-142 | GACAGUGCAGUCACCCAUAAAGUAGAAAGCACUACUAACAGCACUGGAGGGUGUAGUGUUUCCUACUUUAUGGAUGAGUGUACUGUG | 26 | ACAGCACUGGAGGG | 43 | CAUAAAGUAGAAAGCACUACU | 14 | UGUAGUGUUUCCUACUUUAUGGA | 22 |
| hsa-mir-143 | GCGCAGCGCCCUGUCUCCCAGCCUGAGGUGCAGUGCUGCAUCUCUGGUCAGUUGGGAGUCUGAGAUGAAGCACUGUAGCUCAGGAAGAGAGAAGUUGUUCUGCAGC | 33 | CAGUUGGGAGUC | 42 | GGUGCAGUGCUGCAUCUCUGGU | 36 | UGAGAUGAAGCACUGUAGCUC | 29 |
| hsa-mir-144 | UGGGGCCCUGGCUGGGAUAUCAUCAUAUACUGUAAGUUUGCGAUGAGACACUACAGUAUAGAUGAUGUACUAGUCCGGGCACCCCC | 27 | UUUGCGAUGAGACAC | 27 | GGAUAUCAUCAUAUACUGUAAG | 18 | UACAGUAUAGAUGAUGUACU | 20 |
| hsa-mir-145 | CACCUUGUCCUCACGGUCCAGUUUUCCCAGGAAUCCCUUAGAUGCUAAGAUGGGGAUUCCUGGAAAUACUGUUCUUGAGGUCAUGGUU | 24 | UAGAUGCUAAGAUGG | 33 | GUCCAGUUUUCCCAGGAAUCCCU | 17 | GGAUUCCUGGAAAUACUGUUCU | 23 |
| hsa-mir-146a | CCGAUGUGUAUCCUCAGCUUUGAGAACUGAAUUCCAUGGGUUGUGUCAGUGUCAGACCUCUGAAAUUCAGUUCUUCAGCUGGGAUAUCUCUGUCAUCGU | 23 | GUGUCAGUGUCAGA | 36 | UGAGAACUGAAUUCCAUGGGUU | 27 | CCUCUGAAAUUCAGUUCUUCAG | 14 |
| hsa-mir-146b | CCUGGCACUGAGAACUGAAUUCCAUAGGCUGUGAGCUCUAGCAAUGCCCUGUGGACUCAGUUCUGGUGCCCGG | 29 | GUGAGCUCUAGCAA | 29 | UGAGAACUGAAUUCCAUAGGCU | 23 | UGCCCUGUGGACUCAGUUCUGG | 32 |
| hsa-mir-148a | GAGGCAAAGUUCUGAGACACUCCGACUCUGAGUAUGAUAGAAGUCAGUGCACUACAGAACUUUGUCUC | 24 | CUGAGUAUGAUAGAAG | 31 | AAAGUUCUGAGACACUCCGACU | 18 | UCAGUGCACUACAGAACUUUGU | 18 |
| hsa-mir-148b | CAAGCACGAUUAGCAUUUGAGGUGAAGUUCUGUUAUACACUCAGGCUGUGGCUCUCUGAAAGUCAGUGCAUCACAGAACUUUGUCUCGAAAGCUUUCUA | 22 | UGUGGCUCUCUGAAAG | 31 | AAGUUCUGUUAUACACUCAGGC | 18 | UCAGUGCAUCACAGAACUUUGU | 18 |
| hsa-mir-149 | GCCGGCGCCCGAGCUCUGGCUCCGUGUCUUCACUCCCGUGCUUGUCCGAGGAGGGAGGGAGGGACGGGGGCUGUGCUGGGGCAGCUGGA | 44 | GUGCUUGUCCGAGGAGGG | 50 | UCUGGCUCCGUGUCUUCACUCCC | 17 | AGGGAGGGACGGGGGCUGUGC | 62 |
| hsa-mir-150 | CUCCCCAUGGCCCUGUCUCCCAACCCUUGUACCAGUGCUGGGCUCAGACCCUGGUACAGGCCUGGGGGACAGGGACCUGGGGAC | 31 | CUGGGCUCAGACC | 31 | UCUCCCAACCCUUGUACCAGUG | 14 | CUGGUACAGGCCUGGGGGACAG | 45 |
| hsa-mir-151a | UUUCCUGCCCUCGAGGAGCUCACAGUCUAGUAUGUCUCAUCCCCUACUAGACUGAAGCUCCUUGAGGACAGGGAUGGUCAUACUCACCUC | 21 | AUGUCUCAUCCCCUA | 7 | UCGAGGAGCUCACAGUCUAGU | 29 | CUAGACUGAAGCUCCUUGAGG | 29 |
| hsa-mir-152 | UGUCCCCCCCGGCCCAGGUUCUGUGAUACACUCCGACUCGGGCUCUGGAGCAGUCAGUGCAUGACAGAACUUGGGCCCGGAAGGACC | 30 | CGGGCUCUGGAGCAG | 47 | AGGUUCUGUGAUACACUCCGACU | 22 | UCAGUGCAUGACAGAACUUGG | 29 |
| hsa-mir-153-2 | AGCGGUGGCCAGUGUCAUUUUUGUGAUGUUGCAGCUAGUAAUAUGAGCCCAGUUGCAUAGUCACAAAAGUGAUCAUUGGAAACUGUG | 28 | AGUAAUAUGAGCCCAG | 25 | UCAUUUUUGUGAUGUUGCAGCU | 23 | UUGCAUAGUCACAAAAGUGAUC | 18 |
| hsa-mir-154 | GUGGUACUUGAAGAUAGGUUAUCCGUGUUGCCUUCGCUUUAUUUGUGACGAAUCAUACACGGUUGACCUAUUUUUCAGUACCAA | 21 | CUUUAUUUGUGACG | 21 | UAGGUUAUCCGUGUUGCCUUCG | 27 | AAUCAUACACGGUUGACCUAUU | 14 |
| hsa-mir-155 | CUGUUAAUGCUAAUCGUGAUAGGGGUUUUUGCCUCCAACUGACUCCUACAUAUUAGCAUUAACAG | 18 | UUUUGCCUCCAACUGA | 13 | UUAAUGCUAAUCGUGAUAGGGGU | 30 | CUCCUACAUAUUAGCAUUAACA | 5 |
| hsa-mir-15a | CCUUGGAGUAAAGUAGCAGCACAUAAUGGUUUGUGGAUUUUGAAAAGGUGCAGGCCAUAUUGUGCUGCCUCAAAAAUACAAGG | 27 | GAUUUUGAAAAGGUG | 33 | UAGCAGCACAUAAUGGUUUGUG | 27 | CAGGCCAUAUUGUGCUGCCUCA | 23 |
| hsa-mir-15b | UUGAGGCCUUAAAGUACUGUAGCAGCACAUCAUGGUUUACAUGCUACAGUCAAGAUGCGAAUCAUUAUUUGCUGCUCUAGAAAUUUAAGGAAAUUCAU | 19 | UGCUACAGUCAAGAUG | 25 | UAGCAGCACAUCAUGGUUUACA | 18 | CGAAUCAUUAUUUGCUGCUCUA | 14 |
| hsa-mir-16-1 | GUCAGCAGUGCCUUAGCAGCACGUAAAUAUUGGCGUUAAGAUUCUAAAAUUAUCUCCAGUAUUAACUGUGCUGCUGAAGUAAGGUUGAC | 22 | UUAAGAUUCUAAAAUUAUCU | 5 | UAGCAGCACGUAAAUAUUGGCG | 27 | CCAGUAUUAACUGUGCUGCUGA | 23 |
| hsa-mir-16-2 | GUUCCACUCUAGCAGCACGUAAAUAUUGGCGUAGUGAAAUAUAUAUUAAACACCAAUAUUACUGUGCUGCUUUAGUGUGAC | 19 | UAGUGAAAUAUAUAUUAAACA | 10 | UAGCAGCACGUAAAUAUUGGCG | 27 | CCAAUAUUACUGUGCUGCUUUA | 14 |
| hsa-mir-17 | GUCAGAAUAAUGUCAAAGUGCUUACAGUGCAGGUAGUGAUAUGUGCAUCUACUGCAGUGAAGGCACUUGUAGCAUUAUGGUGAC | 27 | UGAUAUGUGCAUCU | 21 | CAAAGUGCUUACAGUGCAGGUAG | 30 | ACUGCAGUGAAGGCACUUGUAG | 32 |
| hsa-mir-181a-1 | UGAGUUUUGAGGUUGCUUCAGUGAACAUUCAACGCUGUCGGUGAGUUUGGAAUUAAAAUCAAAACCAUCGACCGUUGAUUGUACCCUAUGGCUAACCAUCAUCUACUCCA | 20 | UUGGAAUUAAAAUCAAA | 12 | AACAUUCAACGCUGUCGGUGAGU | 26 | ACCAUCGACCGUUGAUUGUACC | 18 |
| hsa-mir-181a-2 | AGAAGGGCUAUCAGGCCAGCCUUCAGAGGACUCCAAGGAACAUUCAACGCUGUCGGUGAGUUUGGGAUUUGAAAAAACCACUGACCGUUGACUGUACCUUGGGGUCCUUA | 27 | UUGGGAUUUGAAAAA | 27 | AACAUUCAACGCUGUCGGUGAGU | 26 | ACCACUGACCGUUGACUGUACC | 18 |
| hsa-mir-181b-1 | CCUGUGCAGAGAUUAUUUUUUAAAAGGUCACAAUCAACAUUCAUUGCUGUCGGUGGGUUGAACUGUGUGGACAAGCUCACUGAACAAUGAAUGCAACUGUGGCCCCGCUU | 24 | UGAACUGUGUGGACAAG | 35 | AACAUUCAUUGCUGUCGGUGGGU | 30 | CUCACUGAACAAUGAAUGCAA | 14 |
| hsa-mir-181b-2 | CUGAUGGCUGCACUCAACAUUCAUUGCUGUCGGUGGGUUUGAGUCUGAAUCAACUCACUGAUCAAUGAAUGCAAACUGCGGACCAAACA | 22 | UUGAGUCUGAAUCAA | 20 | AACAUUCAUUGCUGUCGGUGGGU | 30 | CUCACUGAUCAAUGAAUGCA | 15 |
| hsa-mir-181c | CGGAAAAUUUGCCAAGGGUUUGGGGGAACAUUCAACCUGUCGGUGAGUUUGGGCAGCUCAGGCAAACCAUCGACCGUUGAGUGGACCCUGAGGCCUGGAAUUGCCAUCCU | 31 | UUGGGCAGCUCAGGCA | 38 | AACAUUCAACCUGUCGGUGAGU | 23 | AACCAUCGACCGUUGAGUGGAC | 27 |
| hsa-mir-181d | GUCCCCUCCCCUAGGCCACAGCCGAGGUCACAAUCAACAUUCAUUGUUGUCGGUGGGUUGUGAGGACUGAGGCCAGACCCACCGGGGGAUGAAUGUCACUGUGGCUGGGCCAGACACGGCUUAAGGGGAAUGGGGAC | 34 | UGUGAGGACUGAGGCCAGAC | 40 | AACAUUCAUUGUUGUCGGUGGGU | 30 | CCACCGGGGGAUGAAUGUCAC | 33 |
| hsa-mir-182 | GAGCUGCUUGCCUCCCCCCGUUUUUGGCAAUGGUAGAACUCACACUGGUGAGGUAACAGGAUCCGGUGGUUCUAGACUUGCCAACUAUGGGGCGAGGACUCAGCCGGCAC | 30 | GGUGAGGUAACAGGAUCCGG | 45 | UUUGGCAAUGGUAGAACUCACACU | 21 | UGGUUCUAGACUUGCCAACUA | 19 |
| hsa-mir-183 | CCGCAGAGUGUGACUCCUGUUCUGUGUAUGGCACUGGUAGAAUUCACUGUGAACAGUCUCAGUCAGUGAAUUACCGAAGGGCCAUAAACAGAGCAGAGACAGAUCCACGA | 26 | GUGAACAGUCUCAGUCA | 24 | UAUGGCACUGGUAGAAUUCACU | 23 | GUGAAUUACCGAAGGGCCAUAA | 27 |
| hsa-mir-185 | AGGGGGCGAGGGAUUGGAGAGAAAGGCAGUUCCUGAUGGUCCCCUCCCCAGGGGCUGGCUUUCCUCUGGUCCUUCCCUCCCA | 33 | UGGUCCCCUCCCC | 15 | UGGAGAGAAAGGCAGUUCCUGA | 36 | AGGGGCUGGCUUUCCUCUGGUC | 36 |
| hsa-mir-186 | UGCUUGUAACUUUCCAAAGAAUUCUCCUUUUGGGCUUUCUGGUUUUAUUUUAAGCCCAAAGGUGAAUUUUUUGGGAAGUUUGAGCU | 21 | UUCUGGUUUUAUUUUAA | 12 | CAAAGAAUUCUCCUUUUGGGCU | 18 | GCCCAAAGGUGAAUUUUUUGGG | 32 |
| hsa-mir-187 | GGUCGGGCUCACCAUGACACAGUGUGAGACCUCGGGCUACAACACAGGACCCGGGCGCUGCUCUGACCCCUCGUGUCUUGUGUUGCAGCCGGAGGGACGCAGGUCCGCA | 33 | GCUGCUCUGACCCC | 21 | GGCUACAACACAGGACCCGGGC | 32 | UCGUGUCUUGUGUUGCAGCCGG | 36 |
| hsa-mir-188 | UGCUCCCUCUCUCACAUCCCUUGCAUGGUGGAGGGUGAGCUUUCUGAAAACCCCUCCCACAUGCAGGGUUUGCAGGAUGGCGAGCC | 27 | UGAGCUUUCUGAAAACCC | 17 | CAUCCCUUGCAUGGUGGAGGG | 38 | CUCCCACAUGCAGGGUUUGCA | 24 |
| hsa-mir-18a | UGUUCUAAGGUGCAUCUAGUGCAGAUAGUGAAGUAGAUUAGCAUCUACUGCCCUAAGUGCUCCUUCUGGCA | 24 | UGAAGUAGAUUAGCAUCU | 22 | UAAGGUGCAUCUAGUGCAGAUAG | 30 | ACUGCCCUAAGUGCUCCUUCUGG | 22 |
| hsa-mir-18b | UGUGUUAAGGUGCAUCUAGUGCAGUUAGUGAAGCAGCUUAGAAUCUACUGCCCUAAAUGCCCCUUCUGGCA | 24 | UGAAGCAGCUUAGAAUCUAC | 20 | UAAGGUGCAUCUAGUGCAGUUAG | 30 | UGCCCUAAAUGCCCCUUCUGGC | 18 |
| hsa-mir-190a | UGCAGGCCUCUGUGUGAUAUGUUUGAUAUAUUAGGUUGUUAUUUAAUCCAACUAUAUAUCAAACAUAUUCCUACAGUGUCUUGCC | 16 | UGUUAUUUAAUCCAA | 7 | UGAUAUGUUUGAUAUAUUAGGU | 23 | CUAUAUAUCAAACAUAUUCCU | 0 |
| hsa-mir-191 | CGGCUGGACAGCGGGCAACGGAAUCCCAAAAGCAGCUGUUGUCUCCAGAGCAUUCCAGCUGCGCUUGGAUUUCGUCCCCUGCUCUCCUGCCU | 26 | UUGUCUCCAGAGCAUUCCA | 16 | CAACGGAAUCCCAAAAGCAGCUG | 22 | GCUGCGCUUGGAUUUCGUCCCC | 27 |
| hsa-mir-192 | GCCGAGACCGAGUGCACAGGGCUCUGACCUAUGAAUUGACAGCCAGUGCUCUCGUCUCCCCUCUGGCUGCCAAUUCCAUAGGUCACAGGUAUGUUCGCCUCAAUGCCAGC | 25 | AGUGCUCUCGUCUCCCCUCUGG | 23 | CUGACCUAUGAAUUGACAGCC | 19 | CUGCCAAUUCCAUAGGUCACAG | 18 |
| hsa-mir-193a | CGAGGAUGGGAGCUGAGGGCUGGGUCUUUGCGGGCGAGAUGAGGGUGUCGGAUCAACUGGCCUACAAAGUCCCAGUUCUCGGCCCCCG | 39 | GGGUGUCGGAUC | 50 | UGGGUCUUUGCGGGCGAGAUGA | 45 | AACUGGCCUACAAAGUCCCAGU | 18 |
| hsa-mir-193b | GUGGUCUCAGAAUCGGGGUUUUGAGGGCGAGAUGAGUUUAUGUUUUAUCCAACUGGCCCUCAAAGUCCCGCUUUUGGGGUCAU | 30 | GUUUAUGUUUUAUCC | 13 | CGGGGUUUUGAGGGCGAGAUGA | 50 | AACUGGCCCUCAAAGUCCCGCU | 18 |
| hsa-mir-194-2 | UGGUUCCCGCCCCCUGUAACAGCAACUCCAUGUGGAAGUGCCCACUGGUUCCAGUGGGGCUGCUGUUAUCUGGGGCGAGGGCCAG | 33 | AGUGCCCACUGGUU | 29 | UGUAACAGCAACUCCAUGUGGA | 23 | CCAGUGGGGCUGCUGUUAUCUG | 36 |
| hsa-mir-195 | AGCUUCCCUGGCUCUAGCAGCACAGAAAUAUUGGCACAGGGAAGCGAGUCUGCCAAUAUUGGCUGUGCUGCUCCAGGCAGGGUGGUG | 32 | ACAGGGAAGCGAGUCUG | 41 | UAGCAGCACAGAAAUAUUGGC | 24 | CCAAUAUUGGCUGUGCUGCUCC | 23 |
| hsa-mir-196a-2 | UGCUCGCUCAGCUGAUCUGUGGCUUAGGUAGUUUCAUGUUGUUGGGAUUGAGUUUUGAACUCGGCAACAAGAAACUGCCUGAGUUACAUCAGUCGGUUUUCGUCGAGGGC | 29 | AUUGAGUUUUGAACU | 20 | UAGGUAGUUUCAUGUUGUUGGG | 36 | CGGCAACAAGAAACUGCCUGAG | 27 |
| hsa-mir-196b | ACUGGUCGGUGAUUUAGGUAGUUUCCUGUUGUUGGGAUCCACCUUUCUCUCGACAGCACGACACUGCCUUCAUUACUUCAGUUG | 23 | UCCACCUUUCUC | 0 | UAGGUAGUUUCCUGUUGUUGGG | 36 | UCGACAGCACGACACUGCCUUC | 18 |
| hsa-mir-197 | GGCUGUGCCGGGUAGAGAGGGCAGUGGGAGGUAAGAGCUCUUCACCCUUCACCACCUUCUCCACCCAGCAUGGCC | 31 | UAAGAGCUCUUCACCC | 13 | CGGGUAGAGAGGGCAGUGGGAGG | 61 | UUCACCACCUUCUCCACCCAGC | 5 |
| hsa-mir-199a-1 | GCCAACCCAGUGUUCAGACUACCUGUUCAGGAGGCUCUCAAUGUGUACAGUAGUCUGCACAUUGGUUAGGC | 25 | AGGAGGCUCUCAAUGUGU | 33 | CCCAGUGUUCAGACUACCUGUUC | 17 | ACAGUAGUCUGCACAUUGGUUA | 23 |
| hsa-mir-199a-2 | AGGAAGCUUCUGGAGAUCCUGCUCCGUCGCCCCAGUGUUCAGACUACCUGUUCAGGACAAUGCCGUUGUACAGUAGUCUGCACAUUGGUUAGACUGGGCAAGGGAGAGCA | 29 | AGGACAAUGCCGUUGU | 31 | CCCAGUGUUCAGACUACCUGUUC | 17 | ACAGUAGUCUGCACAUUGGUUA | 23 |
| hsa-mir-199b | CCAGAGGACACCUCCACUCCGUCUACCCAGUGUUUAGACUAUCUGUUCAGGACUCCCAAAUUGUACAGUAGUCUGCACAUUGGUUAGGCUGGGCUGGGUUAGACCCUCGG | 25 | AGGACUCCCAAAUUGU | 19 | CCCAGUGUUUAGACUAUCUGUUC | 17 | ACAGUAGUCUGCACAUUGGUUA | 23 |
| hsa-mir-19a | GCAGUCCUCUGUUAGUUUUGCAUAGUUGCACUACAAGAAGAAUGUAGUUGUGCAAAUCUAUGCAAAACUGAUGGUGGCCUGC | 24 | AGAAGAAUGUAGU | 31 | AGUUUUGCAUAGUUGCACUACA | 18 | UGUGCAAAUCUAUGCAAAACUGA | 17 |
| hsa-mir-19b-1 | CACUGUUCUAUGGUUAGUUUUGCAGGUUUGCAUCCAGCUGUGUGAUAUUCUGCUGUGCAAAUCCAUGCAAAACUGACUGUGGUAGUG | 25 | UGUGUGAUAUUCUGC | 27 | AGUUUUGCAGGUUUGCAUCCAGC | 26 | UGUGCAAAUCCAUGCAAAACUGA | 17 |
| hsa-mir-19b-2 | ACAUUGCUACUUACAAUUAGUUUUGCAGGUUUGCAUUUCAGCGUAUAUAUGUAUAUGUGGCUGUGCAAAUCCAUGCAAAACUGAUUGUGAUAAUGU | 20 | GCGUAUAUAUGUAUAUGUGGC | 29 | AGUUUUGCAGGUUUGCAUUUCA | 23 | UGUGCAAAUCCAUGCAAAACUGA | 17 |
| hsa-mir-200a | CCGGGCCCCUGUGAGCAUCUUACCGGACAGUGCUGGAUUUCCCAGCUUGACUCUAACACUGUCUGGUAACGAUGUUCAAAGGUGACCCGC | 26 | UUUCCCAGCUUGACUC | 13 | CAUCUUACCGGACAGUGCUGGA | 27 | UAACACUGUCUGGUAACGAUGU | 23 |
| hsa-mir-200b | CCAGCUCGGGCAGCCGUGGCCAUCUUACUGGGCAGCAUUGGAUGGAGUCAGGUCUCUAAUACUGCCUGGUAAUGAUGACGGCGGAGCCCUGCACG | 33 | UGGAGUCAGGUCUC | 36 | CAUCUUACUGGGCAGCAUUGGA | 27 | UAAUACUGCCUGGUAAUGAUGA | 23 |
| hsa-mir-200c | CCCUCGUCUUACCCAGCAGUGUUUGGGUGCGGUUGGGAGUCUCUAAUACUGCCGGGUAAUGAUGGAGG | 34 | GUGCGGUUGGGAGUCUC | 47 | CGUCUUACCCAGCAGUGUUUGG | 27 | UAAUACUGCCGGGUAAUGAUGGA | 30 |
| hsa-mir-202 | CGCCUCAGAGCCGCCCGCCGUUCCUUUUUCCUAUGCAUAUACUUCUUUGAGGAUCUGGCCUAAAGAGGUAUAGGGCAUGGGAAAACGGGGCGGUCGGGUCCUCCCCAGCG | 29 | AGGAUCUGGCCUAA | 29 | UUCCUAUGCAUAUACUUCUUUG | 9 | AGAGGUAUAGGGCAUGGGAA | 45 |
| hsa-mir-203a | GUGUUGGGGACUCGCGCGCUGGGUCCAGUGGUUCUUAACAGUUCAACAGUUCUGUAGCGCAAUUGUGAAAUGUUUAGGACCACUAGACCCGGCGGGCGCGGCGACAGCGA | 34 | CUGUAGCGCAAUU | 23 | AGUGGUUCUUAACAGUUCAACAGUU | 20 | UGAAAUGUUUAGGACCACUAG | 24 |
| hsa-mir-203b | GCGCCCGCCGGGUCUAGUGGUCCUAAACAUUUCACAAUUGCGCUACAGAACUGUUGAACUGUUAAGAACCACUGGACCCAGCGCGC | 24 | AUUGCGCUACAGAACUG | 24 | UAGUGGUCCUAAACAUUUCACA | 14 | UUGAACUGUUAAGAACCACUGGA | 22 |
| hsa-mir-204 | GGCUACAGUCUUUCUUCAUGUGACUCGUGGACUUCCCUUUGUCAUCCUAUGCCUGAGAAUAUAUGAAGGAGGCUGGGAAGGCAAAGGGACGUUCAAUUGUCAUCACUGGC | 26 | GAGAAUAUAUGAAGGAG | 35 | UUCCCUUUGUCAUCCUAUGCCU | 9 | GCUGGGAAGGCAAAGGGACGU | 48 |
| hsa-mir-205 | AAAGAUCCUCAGACAAUCCAUGUGCUUCUCUUGUCCUUCAUUCCACCGGAGUCUGUCUCAUACCCAACCAGAUUUCAGUGGAGUGAAGUUCAGGAGGCAUGGAGCUGACA | 22 | UCUCAUACCCAACCA | 0 | UCCUUCAUUCCACCGGAGUCUG | 18 | GAUUUCAGUGGAGUGAAGUUC | 33 |
| hsa-mir-208a | UGACGGGCGAGCUUUUGGCCCGGGUUAUACCUGAUGCUCACGUAUAAGACGAGCAAAAAGCUUGUUGGUCA | 30 | CUGAUGCUCACGU | 23 | GAGCUUUUGGCCCGGGUUAUAC | 32 | AUAAGACGAGCAAAAAGCUUGU | 23 |
| hsa-mir-208b | CCUCUCAGGGAAGCUUUUUGCUCGAAUUAUGUUUCUGAUCCGAAUAUAAGACGAACAAAAGGUUUGUCUGAGGGCAG | 25 | UUCUGAUCCGAAU | 15 | AAGCUUUUUGCUCGAAUUAUGU | 18 | AUAAGACGAACAAAAGGUUUGU | 23 |
| hsa-mir-20a | GUAGCACUAAAGUGCUUAUAGUGCAGGUAGUGUUUAGUUAUCUACUGCAUUAUGAGCACUUAAAGUACUGC | 23 | UGUUUAGUUAUCU | 15 | UAAAGUGCUUAUAGUGCAGGUAG | 30 | ACUGCAUUAUGAGCACUUAAAG | 18 |
| hsa-mir-20b | AGUACCAAAGUGCUCAUAGUGCAGGUAGUUUUGGCAUGACUCUACUGUAGUAUGGGCACUUCCAGUACU | 25 | UUUUGGCAUGACUCU | 20 | CAAAGUGCUCAUAGUGCAGGUAG | 30 | ACUGUAGUAUGGGCACUUCCAG | 27 |
| hsa-mir-21 | UGUCGGGUAGCUUAUCAGACUGAUGUUGACUGUUGAAUCUCAUGGCAACACCAGUCGAUGGGCUGUCUGACA | 28 | CUGUUGAAUCUCAUGG | 25 | UAGCUUAUCAGACUGAUGUUGA | 23 | CAACACCAGUCGAUGGGCUGU | 29 |
| hsa-mir-210 | ACCCGGCAGUGCCUCCAGGCGCAGGGCAGCCCCUGCCCACCGCACACUGCGCUGCCCCAGACCCACUGUGCGUGUGACAGCGGCUGAUCUGUGCCUGGGCAGCGCGACCC | 31 | CGCUGCCCCAGACCCA | 19 | AGCCCCUGCCCACCGCACACUG | 18 | CUGUGCGUGUGACAGCGGCUGA | 41 |
| hsa-mir-211 | UCACCUGGCCAUGUGACUUGUGGGCUUCCCUUUGUCAUCCUUCGCCUAGGGCUCUGAGCAGGGCAGGGACAGCAAAGGGGUGCUCAGUUGUCACUUCCCACAGCACGGAG | 30 | AGGGCUCUGAGCAGG | 47 | UUCCCUUUGUCAUCCUUCGCCU | 9 | GCAGGGACAGCAAAGGGGUGC | 48 |
| hsa-mir-212 | CGGGGCACCCCGCCCGGACAGCGCGCCGGCACCUUGGCUCUAGACUGCUUACUGCCCGGGCCGCCCUCAGUAACAGUCUCCAGUCACGGCCACCGACGCCUGGCCCCGCC | 28 | GCCCGGGCCGCCCUCAG | 35 | ACCUUGGCUCUAGACUGCUUACU | 17 | UAACAGUCUCCAGUCACGGCC | 19 |
| hsa-mir-214 | GGCCUGGCUGGACAGAGUUGUCAUGUGUCUGCCUGUCUACACUUGCUGUGCAGAACAUCCGCUCACCUGUACAGCAGGCACAGACAGGCAGUCACAUGACAACCCAGCCU | 25 | AGAACAUCCGCUCACCUGU | 16 | UGCCUGUCUACACUUGCUGUGC | 23 | ACAGCAGGCACAGACAGGCAGU | 32 |
| hsa-mir-215 | AUCAUUCAGAAAUGGUAUACAGGAAAAUGACCUAUGAAUUGACAGACAAUAUAGCUGAGUUUGUCUGUCAUUUCUUUAGGCCAAUAUUCUGUAUGACUGUGCUACUUCAA | 18 | AAUAUAGCUGAGUUUG | 25 | AUGACCUAUGAAUUGACAGAC | 19 | UCUGUCAUUUCUUUAGGCCAAUA | 13 |
| hsa-mir-216a | GAUGGCUGUGAGUUGGCUUAAUCUCAGCUGGCAACUGUGAGAUGUUCAUACAAUCCCUCACAGUGGUCUCUGGGAUUAUGCUAAACAGAGCAAUUUCCUAGCCCUCACGA | 24 | GAUGUUCAUACAAUCCC | 12 | UAAUCUCAGCUGGCAACUGUGA | 23 | UCACAGUGGUCUCUGGGAUUAU | 27 |
| hsa-mir-216b | GCAGACUGGAAAAUCUCUGCAGGCAAAUGUGAUGUCACUGAGGAAAUCACACACUUACCCGUAGAGAUUCUACAGUCUGACA | 22 | UGUCACUGAGGAAAUC | 25 | AAAUCUCUGCAGGCAAAUGUGA | 23 | ACACACUUACCCGUAGAGAUUCUA | 13 |
| hsa-mir-218-1 | GUGAUAAUGUAGCGAGAUUUUCUGUUGUGCUUGAUCUAACCAUGUGGUUGCGAGGUAUGAGUAAAACAUGGUUCCGUCAAGCACCAUGGAACGUCACGCAGCUUUCUACA | 25 | GGUUGCGAGGUAUGAGUAAAAC | 36 | UUGUGCUUGAUCUAACCAUGU | 19 | AUGGUUCCGUCAAGCACCAUGG | 27 |
| hsa-mir-218-2 | GACCAGUCGCUGCGGGGCUUUCCUUUGUGCUUGAUCUAACCAUGUGGUGGAACGAUGGAAACGGAACAUGGUUCUGUCAAGCACCGCGGAAAGCACCGUGCUCUCCUGCA | 29 | GGUGGAACGAUGGAAACGGAA | 43 | UUGUGCUUGAUCUAACCAUGU | 19 | CAUGGUUCUGUCAAGCACCGCG | 27 |
| hsa-mir-219a-1 | CCGCCCCGGGCCGCGGCUCCUGAUUGUCCAAACGCAAUUCUCGAGUCUAUGGCUCCGGCCGAGAGUUGAGUCUGGACGUCCCGAGCCGCCGCCCCCAAACCUCGAGCGGG | 30 | CGAGUCUAUGGCUCCGGCCG | 35 | UGAUUGUCCAAACGCAAUUCU | 14 | AGAGUUGAGUCUGGACGUCCCG | 36 |
| hsa-mir-219a-2 | ACUCAGGGGCUUCGCCACUGAUUGUCCAAACGCAAUUCUUGUACGAGUCUGCGGCCAACCGAGAAUUGUGGCUGGACAUCUGUGGCUGAGCUCCGGG | 30 | UGUACGAGUCUGCGGCCAACCG | 32 | UGAUUGUCCAAACGCAAUUCU | 14 | AGAAUUGUGGCUGGACAUCUGU | 32 |
| hsa-mir-219b | GGAGCUCAGCCACAGAUGUCCAGCCACAAUUCUCGGUUGGCCGCAGACUCGUACAAGAAUUGCGUUUGGACAAUCAGUGGCGAAGCCC | 27 | GUUGGCCGCAGACUCGUACA | 30 | AGAUGUCCAGCCACAAUUCUCG | 18 | AGAAUUGCGUUUGGACAAUCAGU | 26 |
| hsa-mir-22 | GGCUGAGCCGCAGUAGUUCUUCAGUGGCAAGCUUUAUGUCCUGACCCAGCUAAAGCUGCCAGUUGAAGAACUGUUGCCCUCUGCC | 26 | UGUCCUGACCCAGCUA | 19 | AGUUCUUCAGUGGCAAGCUUUA | 23 | AAGCUGCCAGUUGAAGAACUGU | 27 |
| hsa-mir-221 | UGAACAUCCAGGUCUGGGGCAUGAACCUGGCAUACAAUGUAGAUUUCUGUGUUCGUUAGGCAACAGCUACAUUGUCUGCUGGGUUUCAGGCUACCUGGAAACAUGUUCUC | 25 | CUGUGUUCGUUAGGCAAC | 28 | ACCUGGCAUACAAUGUAGAUUU | 18 | AGCUACAUUGUCUGCUGGGUUUC | 26 |
| hsa-mir-222 | GCUGCUGGAAGGUGUAGGUACCCUCAAUGGCUCAGUAGCCAGUGUAGAUCCUGUCUUUCGUAAUCAGCAGCUACAUCUGGCUACUGGGUCUCUGAUGGCAUCUUCUAGCU | 26 | GUCUUUCGUAAUCAGC | 19 | CUCAGUAGCCAGUGUAGAUCCU | 23 | AGCUACAUCUGGCUACUGGGU | 29 |
| hsa-mir-223 | CCUGGCCUCCUGCAGUGCCACGCUCCGUGUAUUUGACAAGCUGAGUUGGACACUCCAUGUGGUAGAGUGUCAGUUUGUCAAAUACCCCAAGUGCGGCACAUGCUUACCAG | 25 | GGACACUCCAUGUGGUAGAG | 35 | CGUGUAUUUGACAAGCUGAGUU | 27 | UGUCAGUUUGUCAAAUACCCCA | 14 |
| hsa-mir-224 | GGGCUUUCAAGUCACUAGUGGUUCCGUUUAGUAGAUGAUUGUGCAUUGUUUCAAAAUGGUGCCCUAGUGACUACAAAGCCC | 25 | UAGUAGAUGAUUGUGCAUUGUUUC | 25 | CAAGUCACUAGUGGUUCCGUU | 24 | AAAAUGGUGCCCUAGUGACUACA | 22 |
| hsa-mir-23a | GGCCGGCUGGGGUUCCUGGGGAUGGGAUUUGCUUCCUGUCACAAAUCACAUUGCCAGGGAUUUCCAACCGACC | 30 | GCUUCCUGUCACAA | 14 | GGGGUUCCUGGGGAUGGGAUUU | 50 | AUCACAUUGCCAGGGAUUUCC | 19 |
| hsa-mir-23b | CUCAGGUGCUCUGGCUGCUUGGGUUCCUGGCAUGCUGAUUUGUGACUUAAGAUUAAAAUCACAUUGCCAGGGAUUACCACGCAACCACGACCUUGGC | 25 | GUGACUUAAGAUUAAA | 19 | UGGGUUCCUGGCAUGCUGAUUU | 32 | AUCACAUUGCCAGGGAUUACC | 19 |
| hsa-mir-24-1 | CUCCGGUGCCUACUGAGCUGAUAUCAGUUCUCAUUUUACACACUGGCUCAGUUCAGCAGGAACAGGAG | 24 | UCUCAUUUUACACAC | 0 | UGCCUACUGAGCUGAUAUCAGU | 23 | UGGCUCAGUUCAGCAGGAACAG | 32 |
| hsa-mir-24-2 | CUCUGCCUCCCGUGCCUACUGAGCUGAAACACAGUUGGUUUGUGUACACUGGCUCAGUUCAGCAGGAACAGGG | 27 | UUGGUUUGUGUACAC | 27 | UGCCUACUGAGCUGAAACACAG | 23 | UGGCUCAGUUCAGCAGGAACAG | 32 |
| hsa-mir-25 | GGCCAGUGUUGAGAGGCGGAGACUUGGGCAAUUGCUGGACGCUGCCCUGGGCAUUGCACUUGUCUCGGUCUGACAGUGCCGGCC | 37 | CUGGACGCUGCCCUGGG | 41 | AGGCGGAGACUUGGGCAAUUG | 43 | CAUUGCACUUGUCUCGGUCUGA | 23 |
| hsa-mir-26a-1 | GUGGCCUCGUUCAAGUAAUCCAGGAUAGGCUGUGCAGGUCCCAAUGGGCCUAUUCUUGGUUACUUGCACGGGGACGC | 31 | GUGCAGGUCCCAAUGGG | 41 | UUCAAGUAAUCCAGGAUAGGCU | 23 | CCUAUUCUUGGUUACUUGCACG | 18 |
| hsa-mir-26a-2 | GGCUGUGGCUGGAUUCAAGUAAUCCAGGAUAGGCUGUUUCCAUCUGUGAGGCCUAUUCUUGAUUACUUGUUUCUGGAGGCAGCU | 29 | GUUUCCAUCUGUGAGG | 31 | UUCAAGUAAUCCAGGAUAGGCU | 23 | CCUAUUCUUGAUUACUUGUUUC | 9 |
| hsa-mir-26b | CCGGGACCCAGUUCAAGUAAUUCAGGAUAGGUUGUGUGCUGUCCAGCCUGUUCUCCAUUACUUGGCUCGGGGACCGG | 30 | UGUGUGCUGUCCAG | 36 | UUCAAGUAAUUCAGGAUAGGU | 24 | CCUGUUCUCCAUUACUUGGCUC | 14 |
| hsa-mir-27a | CUGAGGAGCAGGGCUUAGCUGCUUGUGAGCAGGGUCCACACCAAGUCGUGUUCACAGUGGCUAAGUUCCGCCCCCCAG | 31 | GGGUCCACACCAAGUCGUG | 32 | AGGGCUUAGCUGCUUGUGAGCA | 36 | UUCACAGUGGCUAAGUUCCGC | 24 |
| hsa-mir-27b | ACCUCUCUAACAAGGUGCAGAGCUUAGCUGAUUGGUGAACAGUGAUUGGUUUCCGCUUUGUUCACAGUGGCUAAGUUCUGCACCUGAAGAGAAGGUG | 28 | AGUGAUUGGUUUCCGCUUUG | 30 | AGAGCUUAGCUGAUUGGUGAAC | 32 | UUCACAGUGGCUAAGUUCUGC | 24 |
| hsa-mir-28 | GGUCCUUGCCCUCAAGGAGCUCACAGUCUAUUGAGUUACCUUUCUGACUUUCCCACUAGAUUGUGAGCUCCUGGAGGGCAGGCACU | 24 | UUACCUUUCUGACUUUCC | 6 | AAGGAGCUCACAGUCUAUUGAG | 27 | CACUAGAUUGUGAGCUCCUGGA | 27 |
| hsa-mir-296 | AGGACCCUUCCAGAGGGCCCCCCCUCAAUCCUGUUGUGCCUAAUUCAGAGGGUUGGGUGGAGGCUCUCCUGAAGGGCUCU | 30 | UGUGCCUAAUUCA | 15 | AGGGCCCCCCCUCAAUCCUGU | 19 | GAGGGUUGGGUGGAGGCUCUCC | 50 |
| hsa-mir-299 | AAGAAAUGGUUUACCGUCCCACAUACAUUUUGAAUAUGUAUGUGGGAUGGUAAACCGCUUCUU | 21 | UUUGAAUAUG | 20 | UGGUUUACCGUCCCACAUACAU | 14 | UAUGUGGGAUGGUAAACCGCUU | 32 |
| hsa-mir-29a | AUGACUGAUUUCUUUUGGUGUUCAGAGUCAAUAUAAUUUUCUAGCACCAUCUGAAAUCGGUUAU | 17 | AGUCAAUAUAAUUUUC | 6 | ACUGAUUUCUUUUGGUGUUCAG | 23 | UAGCACCAUCUGAAAUCGGUUA | 18 |
| hsa-mir-29b-1 | CUUCAGGAAGCUGGUUUCAUAUGGUGGUUUAGAUUUAAAUAGUGAUUGUCUAGCACCAUUUGAAAUCAGUGUUCUUGGGGG | 27 | UUUAAAUAGUGAUUGUC | 18 | GCUGGUUUCAUAUGGUGGUUUAGA | 33 | UAGCACCAUUUGAAAUCAGUGUU | 17 |
| hsa-mir-29b-2 | CUUCUGGAAGCUGGUUUCACAUGGUGGCUUAGAUUUUUCCAUCUUUGUAUCUAGCACCAUUUGAAAUCAGUGUUUUAGGAG | 22 | AUUUUUCCAUCUUUGUAUC | 5 | CUGGUUUCACAUGGUGGCUUAG | 32 | UAGCACCAUUUGAAAUCAGUGUU | 17 |
| hsa-mir-29c | AUCUCUUACACAGGCUGACCGAUUUCUCCUGGUGUUCAGAGUCUGUUUUUGUCUAGCACCAUUUGAAAUCGGUUAUGAUGUAGGGGGA | 25 | AGAGUCUGUUUUUGUC | 25 | UGACCGAUUUCUCCUGGUGUUC | 23 | UAGCACCAUUUGAAAUCGGUUA | 18 |
| hsa-mir-301a | ACUGCUAACGAAUGCUCUGACUUUAUUGCACUACUGUACUUUACAGCUAGCAGUGCAAUAGUAUUGUCAAAGCAUCUGAAAGCAGG | 20 | GUACUUUACAGCUAG | 20 | GCUCUGACUUUAUUGCACUACU | 14 | CAGUGCAAUAGUAUUGUCAAAGC | 22 |
| hsa-mir-301b | GCCGCAGGUGCUCUGACGAGGUUGCACUACUGUGCUCUGAGAAGCAGUGCAAUGAUAUUGUCAAAGCAUCUGGGACCA | 29 | GUGCUCUGAGAAG | 38 | GCUCUGACGAGGUUGCACUACU | 27 | CAGUGCAAUGAUAUUGUCAAAGC | 22 |
| hsa-mir-302a | CCACCACUUAAACGUGGAUGUACUUGCUUUGAAACUAAAGAAGUAAGUGCUUCCAUGUUUUGGUGAUGG | 23 | UUGAAACUAAAGAAG | 20 | ACUUAAACGUGGAUGUACUUGCU | 22 | UAAGUGCUUCCAUGUUUUGGUGA | 26 |
| hsa-mir-302b | GCUCCCUUCAACUUUAACAUGGAAGUGCUUUCUGUGACUUUAAAAGUAAGUGCUUCCAUGUUUUAGUAGGAGU | 21 | UGUGACUUUAAAAG | 21 | ACUUUAACAUGGAAGUGCUUUC | 18 | UAAGUGCUUCCAUGUUUUAGUAG | 22 |
| hsa-mir-302c | CCUUUGCUUUAACAUGGGGGUACCUGCUGUGUGAAACAAAAGUAAGUGCUUCCAUGUUUCAGUGGAGG | 28 | UGUGAAACAAAAG | 23 | UUUAACAUGGGGGUACCUGCUG | 32 | UAAGUGCUUCCAUGUUUCAGUGG | 26 |
| hsa-mir-302d | CCUCUACUUUAACAUGGAGGCACUUGCUGUGACAUGACAAAAAUAAGUGCUUCCAUGUUUGAGUGUGG | 24 | UGUGACAUGACAAAAA | 19 | ACUUUAACAUGGAGGCACUUGC | 23 | UAAGUGCUUCCAUGUUUGAGUGU | 26 |
| hsa-mir-30a | GCGACUGUAAACAUCCUCGACUGGAAGCUGUGAAGCCACAGAUGGGCUUUCAGUCGGAUGUUUGCAGCUGC | 30 | CUGUGAAGCCACAGAUGGG | 37 | UGUAAACAUCCUCGACUGGAAG | 23 | CUUUCAGUCGGAUGUUUGCAGC | 27 |
| hsa-mir-30b | ACCAAGUUUCAGUUCAUGUAAACAUCCUACACUCAGCUGUAAUACAUGGAUUGGCUGGGAGGUGGAUGUUUACUUCAGCUGACUUGGA | 24 | GUAAUACAUGGAUUGG | 31 | UGUAAACAUCCUACACUCAGCU | 9 | CUGGGAGGUGGAUGUUUACUUC | 36 |
| hsa-mir-30c-1 | ACCAUGCUGUAGUGUGUGUAAACAUCCUACACUCUCAGCUGUGAGCUCAAGGUGGCUGGGAGAGGGUUGUUUACUCCUUCUGCCAUGGA | 28 | UGUGAGCUCAAGGUGG | 44 | UGUAAACAUCCUACACUCUCAGC | 9 | CUGGGAGAGGGUUGUUUACUCC | 36 |
| hsa-mir-30c-2 | AGAUACUGUAAACAUCCUACACUCUCAGCUGUGGAAAGUAAGAAAGCUGGGAGAAGGCUGUUUACUCUUUCU | 22 | UGUGGAAAGUAAGAAAG | 35 | UGUAAACAUCCUACACUCUCAGC | 9 | CUGGGAGAAGGCUGUUUACUCU | 32 |
| hsa-mir-30d | GUUGUUGUAAACAUCCCCGACUGGAAGCUGUAAGACACAGCUAAGCUUUCAGUCAGAUGUUUGCUGCUAC | 23 | CUGUAAGACACAGCUAAG | 22 | UGUAAACAUCCCCGACUGGAAG | 23 | CUUUCAGUCAGAUGUUUGCUGC | 23 |
| hsa-mir-30e | GGGCAGUCUUUGCUACUGUAAACAUCCUUGACUGGAAGCUGUAAGGUGUUCAGAGGAGCUUUCAGUCGGAUGUUUACAGCGGCAGGCUGCCA | 30 | CUGUAAGGUGUUCAGAGGAG | 40 | UGUAAACAUCCUUGACUGGAAG | 23 | CUUUCAGUCGGAUGUUUACAGC | 23 |
| hsa-mir-31 | GGAGAGGAGGCAAGAUGCUGGCAUAGCUGUUGAACUGGGAACCUGCUAUGCCAACAUAUUGCCAUCUUUCC | 28 | GUUGAACUGGGAACC | 33 | AGGCAAGAUGCUGGCAUAGCU | 33 | UGCUAUGCCAACAUAUUGCCAU | 14 |
| hsa-mir-32 | GGAGAUAUUGCACAUUACUAAGUUGCAUGUUGUCACGGCCUCAAUGCAAUUUAGUGUGUGUGAUAUUUUC | 23 | UGUUGUCACGGCCUCAAUG | 26 | UAUUGCACAUUACUAAGUUGCA | 14 | CAAUUUAGUGUGUGUGAUAUUU | 23 |
| hsa-mir-323a | UUGGUACUUGGAGAGAGGUGGUCCGUGGCGCGUUCGCUUUAUUUAUGGCGCACAUUACACGGUCGACCUCUUUGCAGUAUCUAAUC | 28 | UUUAUUUAUGGCG | 23 | AGGUGGUCCGUGGCGCGUUCGC | 45 | CACAUUACACGGUCGACCUCU | 14 |
| hsa-mir-323b | UGGUACUCGGAGGGAGGUUGUCCGUGGUGAGUUCGCAUUAUUUAAUGAUGCCCAAUACACGGUCGACCUCUUUUCGGUAUCA | 28 | UUAUUUAAUGAUG | 15 | AGGUUGUCCGUGGUGAGUUCGCA | 39 | CCCAAUACACGGUCGACCUCUU | 14 |
| hsa-mir-324 | CUGACUAUGCCUCCCCGCAUCCCCUAGGGCAUUGGUGUAAAGCUGGAGACCCACUGCCCCAGGUGCUGCUGGGGGUUGUAGUC | 30 | AAAGCUGGAGACCC | 29 | CGCAUCCCCUAGGGCAUUGGUGU | 30 | ACUGCCCCAGGUGCUGCUGG | 35 |
| hsa-mir-328 | UGGAGUGGGGGGGCAGGAGGGGCUCAGGGAGAAAGUGCAUACAGCCCCUGGCCCUCUCUGCCCUUCCGUCCCCUG | 37 | AGAAAGUGCAUACAGCCC | 22 | GGGGGGGCAGGAGGGGCUCAGGG | 70 | CUGGCCCUCUCUGCCCUUCCGU | 18 |
| hsa-mir-329-1 | GGUACCUGAAGAGAGGUUUUCUGGGUUUCUGUUUCUUUAAUGAGGACGAAACACACCUGGUUAACCUCUUUUCCAGUAUC | 23 | UUUAAUGAGGACGA | 29 | GAGGUUUUCUGGGUUUCUGUUUC | 30 | AACACACCUGGUUAACCUCUUU | 9 |
| hsa-mir-329-2 | GUGGUACCUGAAGAGAGGUUUUCUGGGUUUCUGUUUCUUUAUUGAGGACGAAACACACCUGGUUAACCUCUUUUCCAGUAUCAA | 23 | UUUAUUGAGGACGA | 29 | GAGGUUUUCUGGGUUUCUGUUUC | 30 | AACACACCUGGUUAACCUCUUU | 9 |
| hsa-mir-330 | CUUUGGCGAUCACUGCCUCUCUGGGCCUGUGUCUUAGGCUCUGCAAGAUCAACCGAGCAAAGCACACGGCCUGCAGAGAGGCAGCGCUCUGCCC | 28 | UCUGCAAGAUCAACCGA | 18 | UCUCUGGGCCUGUGUCUUAGGC | 32 | GCAAAGCACACGGCCUGCAGAGA | 30 |
| hsa-mir-331 | GAGUUUGGUUUUGUUUGGGUUUGUUCUAGGUAUGGUCCCAGGGAUCCCAGAUCAAACCAGGCCCCUGGGCCUAUCCUAGAACCAACCUAAGCUC | 26 | CAGAUCAAACCAG | 15 | CUAGGUAUGGUCCCAGGGAUCC | 32 | GCCCCUGGGCCUAUCCUAGAA | 24 |
| hsa-mir-335 | UGUUUUGAGCGGGGGUCAAGAGCAAUAACGAAAAAUGUUUGUCAUAAACCGUUUUUCAUUAUUGCUCCUGACCUCCUCUCAUUUGCUAUAUUCA | 18 | UUGUCAUAAACCG | 15 | UCAAGAGCAAUAACGAAAAAUGU | 17 | UUUUUCAUUAUUGCUCCUGACC | 9 |
| hsa-mir-337 | GUAGUCAGUAGUUGGGGGGUGGGAACGGCUUCAUACAGGAGUUGAUGCACAGUUAUCCAGCUCCUAUAUGAUGCCUUUCUUCAUCCCCUUCAA | 26 | GAUGCACAGUUAUCCAG | 24 | GAACGGCUUCAUACAGGAGUU | 29 | CUCCUAUAUGAUGCCUUUCUUC | 9 |
| hsa-mir-338 | UCUCCAACAAUAUCCUGGUGCUGAGUGAUGACUCAGGCGACUCCAGCAUCAGUGAUUUUGUUGAAGA | 24 | AUGACUCAGGCGAC | 29 | AACAAUAUCCUGGUGCUGAGUG | 27 | UCCAGCAUCAGUGAUUUUGUUG | 23 |
| hsa-mir-339 | CGGGGCGGCCGCUCUCCCUGUCCUCCAGGAGCUCACGUGUGCCUGCCUGUGAGCGCCUCGACGACAGAGCCGGCGCCUGCCCCAGUGUCUGCGC | 33 | UGUGCCUGCCUG | 33 | UCCCUGUCCUCCAGGAGCUCACG | 22 | UGAGCGCCUCGACGACAGAGCCG | 35 |
| hsa-mir-33a | CUGUGGUGCAUUGUAGUUGCAUUGCAUGUUCUGGUGGUACCCAUGCAAUGUUUCCACAGUGCAUCACAG | 26 | UGUUCUGGUGGUACCCAUG | 32 | GUGCAUUGUAGUUGCAUUGCA | 29 | CAAUGUUUCCACAGUGCAUCAC | 14 |
| hsa-mir-33b | GCGGGCGGCCCCGCGGUGCAUUGCUGUUGCAUUGCACGUGUGUGAGGCGGGUGCAGUGCCUCGGCAGUGCAGCCCGGAGCCGGCCCCUGGCACCAC | 40 | ACGUGUGUGAGGCGGGUG | 56 | GUGCAUUGCUGUUGCAUUGC | 30 | CAGUGCCUCGGCAGUGCAGCCC | 32 |
| hsa-mir-340 | UUGUACCUGGUGUGAUUAUAAAGCAAUGAGACUGAUUGUCAUAUGUCGUUUGUGGGAUCCGUCUCAGUUACUUUAUAGCCAUACCUGGUAUCUUA | 22 | GUCAUAUGUCGUUUGUGGGA | 35 | UUAUAAAGCAAUGAGACUGAUU | 18 | UCCGUCUCAGUUACUUUAUAGC | 14 |
| hsa-mir-342 | GAAACUGGGCUCAAGGUGAGGGGUGCUAUCUGUGAUUGAGGGACAUGGUUAAUGGAAUUGUCUCACACAGAAAUCGCACCCGUCACCUUGGCCUACUUA | 28 | GGGACAUGGUUAAUGGAAUUG | 38 | AGGGGUGCUAUCUGUGAUUGA | 38 | UCUCACACAGAAAUCGCACCCGU | 13 |
| hsa-mir-345 | ACCCAAACCCUAGGUCUGCUGACUCCUAGUCCAGGGCUCGUGAUGGCUGGUGGGCCCUGAACGAGGGGUCUGGAGGCCUGGGUUUGAAUAUCGACAGC | 34 | GUGAUGGCUGGUGG | 57 | GCUGACUCCUAGUCCAGGGCUC | 27 | GCCCUGAACGAGGGGUCUGGAG | 45 |
| hsa-mir-34a | GGCCAGCUGUGAGUGUUUCUUUGGCAGUGUCUUAGCUGGUUGUUGUGAGCAAUAGUAAGGAAGCAAUCAGCAAGUAUACUGCCCUAGAAGUGCUGCACGUUGUGGGGCCC | 32 | UGUGAGCAAUAGUAAGGAAG | 35 | UGGCAGUGUCUUAGCUGGUUGU | 36 | CAAUCAGCAAGUAUACUGCCCU | 14 |
| hsa-mir-34b | GUGCUCGGUUUGUAGGCAGUGUCAUUAGCUGAUUGUACUGUGGUGGUUACAAUCACUAACUCCACUGCCAUCAAAACAAGGCAC | 24 | UACUGUGGUGGUUA | 36 | UAGGCAGUGUCAUUAGCUGAUUG | 30 | CAAUCACUAACUCCACUGCCAU | 5 |
| hsa-mir-34c | AGUCUAGUUACUAGGCAGUGUAGUUAGCUGAUUGCUAAUAGUACCAAUCACUAACCACACGGCCAGGUAAAAAGAUU | 21 | UAAUAGUACC | 10 | AGGCAGUGUAGUUAGCUGAUUGC | 35 | AAUCACUAACCACACGGCCAGG | 18 |
| hsa-mir-361 | GGAGCUUAUCAGAAUCUCCAGGGGUACUUUAUAAUUUCAAAAAGUCCCCCAGGUGUGAUUCUGAUUUGCUUC | 21 | UUUAUAAUUUCAAAAAG | 6 | UUAUCAGAAUCUCCAGGGGUAC | 23 | UCCCCCAGGUGUGAUUCUGAUUU | 22 |
| hsa-mir-362 | CUUGAAUCCUUGGAACCUAGGUGUGAGUGCUAUUUCAGUGCAACACACCUAUUCAAGGAUUCAAA | 20 | GCUAUUUCAGUGC | 23 | AAUCCUUGGAACCUAGGUGUGAGU | 29 | AACACACCUAUUCAAGGAUUCA | 9 |
| hsa-mir-363 | UGUUGUCGGGUGGAUCACGAUGCAAUUUUGAUGAGUAUCAUAGGAGAAAAAUUGCACGGUAUCCAUCUGUAAACC | 25 | UGAUGAGUAUCAUAGGAGAAA | 29 | CGGGUGGAUCACGAUGCAAUUU | 32 | AAUUGCACGGUAUCCAUCUGUA | 18 |
| hsa-mir-365a | ACCGCAGGGAAAAUGAGGGACUUUUGGGGGCAGAUGUGUUUCCAUUCCACUAUCAUAAUGCCCCUAAAAAUCCUUAUUGCUCUUGCA | 22 | UUUCCAUUCCACUAUCA | 0 | AGGGACUUUUGGGGGCAGAUGUG | 48 | UAAUGCCCCUAAAAAUCCUUAU | 5 |
| hsa-mir-365b | AGAGUGUUCAAGGACAGCAAGAAAAAUGAGGGACUUUCAGGGGCAGCUGUGUUUUCUGACUCAGUCAUAAUGCCCCUAAAAAUCCUUAUUGUUCUUGCAGUGUGCAUCGGG | 26 | GUUUUCUGACUCAGUCA | 18 | AGGGACUUUCAGGGGCAGCUGU | 41 | UAAUGCCCCUAAAAAUCCUUAU | 5 |
| hsa-mir-367 | CCAUUACUGUUGCUAAUAUGCAACUCUGUUGAAUAUAAAUUGGAAUUGCACUUUAGCAAUGGUGAUGG | 21 | GUUGAAUAUAAAUUGG | 25 | ACUGUUGCUAAUAUGCAACUCU | 14 | AAUUGCACUUUAGCAAUGGUGA | 23 |
| hsa-mir-369 | UUGAAGGGAGAUCGACCGUGUUAUAUUCGCUUUAUUGACUUCGAAUAAUACAUGGUUGAUCUUUUCUCAG | 21 | UUUAUUGACUUCG | 15 | AGAUCGACCGUGUUAUAUUCGC | 23 | AAUAAUACAUGGUUGAUCUUU | 14 |
| hsa-mir-370 | AGACAGAGAAGCCAGGUCACGUCUCUGCAGUUACACAGCUCACGAGUGCCUGCUGGGGUGGAACCUGGUCUGUCU | 31 | ACAGCUCACGAGU | 23 | CAGGUCACGUCUCUGCAGUUAC | 23 | GCCUGCUGGGGUGGAACCUGGU | 45 |
| hsa-mir-371a | GUGGCACUCAAACUGUGGGGGCACUUUCUGCUCUCUGGUGAAAGUGCCGCCAUCUUUUGAGUGUUAC | 28 | UUCUGCUCUCUGGUGA | 25 | ACUCAAACUGUGGGGGCACU | 30 | AAGUGCCGCCAUCUUUUGAGUGU | 26 |
| hsa-mir-371b | GGUAACACUCAAAAGAUGGCGGCACUUUCACCAGAGAGCAGAAAGUGCCCCCACAGUUUGAGUGCC | 26 | CACCAGAGAGCAGA | 29 | ACUCAAAAGAUGGCGGCACUUU | 23 | AAGUGCCCCCACAGUUUGAGUGC | 26 |
| hsa-mir-372 | GUGGGCCUCAAAUGUGGAGCACUAUUCUGAUGUCCAAGUGGAAAGUGCUGCGACAUUUGAGCGUCAC | 30 | GAUGUCCAAGUGG | 38 | CCUCAAAUGUGGAGCACUAUUCU | 17 | AAAGUGCUGCGACAUUUGAGCGU | 30 |
| hsa-mir-373 | GGGAUACUCAAAAUGGGGGCGCUUUCCUUUUUGUCUGUACUGGGAAGUGCUUCGAUUUUGGGGUGUCCC | 32 | UUUUUGUCUGUACUGG | 25 | ACUCAAAAUGGGGGCGCUUUCC | 27 | GAAGUGCUUCGAUUUUGGGGUGU | 39 |
| hsa-mir-374a | UACAUCGGCCAUUAUAAUACAACCUGAUAAGUGUUAUAGCACUUAUCAGAUUGUAUUGUAAUUGUCUGUGUA | 17 | UUAUAGCA | 13 | UUAUAAUACAACCUGAUAAGUG | 14 | CUUAUCAGAUUGUAUUGUAAUU | 14 |
| hsa-mir-374b | ACUCGGAUGGAUAUAAUACAACCUGCUAAGUGUCCUAGCACUUAGCAGGUUGUAUUAUCAUUGUCCGUGUCU | 21 | UCCUAGCA | 13 | AUAUAAUACAACCUGCUAAGUG | 14 | CUUAGCAGGUUGUAUUAUCAUU | 18 |
| hsa-mir-374c | ACACGGACAAUGAUAAUACAACCUGCUAAGUGCUAGGACACUUAGCAGGUUGUAUUAUAUCCAUCCGAGU | 20 | AGGA | 50 | AUAAUACAACCUGCUAAGUGCU | 14 | CACUUAGCAGGUUGUAUUAUAU | 18 |
| hsa-mir-376a-1 | UAAAAGGUAGAUUCUCCUUCUAUGAGUACAUUAUUUAUGAUUAAUCAUAGAGGAAAAUCCACGUUUUC | 15 | CAUUAUUUAUGAUUA | 7 | GUAGAUUCUCCUUCUAUGAGUA | 18 | AUCAUAGAGGAAAAUCCACGU | 19 |
| hsa-mir-376a-2 | GGUAUUUAAAAGGUAGAUUUUCCUUCUAUGGUUACGUGUUUGAUGGUUAAUCAUAGAGGAAAAUCCACGUUUUCAGUAUC | 21 | UACGUGUUUGAUGGUUA | 29 | GGUAGAUUUUCCUUCUAUGGU | 24 | AUCAUAGAGGAAAAUCCACGU | 19 |
| hsa-mir-376b | CAGUCCUUCUUUGGUAUUUAAAACGUGGAUAUUCCUUCUAUGUUUACGUGAUUCCUGGUUAAUCAUAGAGGAAAAUCCAUGUUUUCAGUAUCAAAUGCUG | 18 | ACGUGAUUCCUGGUUA | 25 | CGUGGAUAUUCCUUCUAUGUUU | 18 | AUCAUAGAGGAAAAUCCAUGUU | 18 |
| hsa-mir-376c | AAAAGGUGGAUAUUCCUUCUAUGUUUAUGUUAUUUAUGGUUAAACAUAGAGGAAAUUCCACGUUUU | 18 | UAUGUUAUUUAUGGUUA | 18 | GGUGGAUAUUCCUUCUAUGUU | 24 | AACAUAGAGGAAAUUCCACGU | 19 |
| hsa-mir-377 | UUGAGCAGAGGUUGCCCUUGGUGAAUUCGCUUUAUUUAUGUUGAAUCACACAAAGGCAACUUUUGUUUG | 23 | GCUUUAUUUAUGUUGA | 19 | AGAGGUUGCCCUUGGUGAAUUC | 32 | AUCACACAAAGGCAACUUUUGU | 14 |
| hsa-mir-378a | AGGGCUCCUGACUCCAGGUCCUGUGUGUUACCUAGAAAUAGCACUGGACUUGGAGUCAGAAGGCCU | 29 | GUUACCUAGAAAUAGC | 19 | CUCCUGACUCCAGGUCCUGUGU | 23 | ACUGGACUUGGAGUCAGAAGGC | 36 |
| hsa-mir-379 | AGAGAUGGUAGACUAUGGAACGUAGGCGUUAUGAUUUCUGACCUAUGUAACAUGGUCCACUAACUCU | 24 | CGUUAUGAUUUCUGACC | 18 | UGGUAGACUAUGGAACGUAGG | 38 | UAUGUAACAUGGUCCACUAACU | 14 |
| hsa-mir-380 | AAGAUGGUUGACCAUAGAACAUGCGCUAUCUCUGUGUCGUAUGUAAUAUGGUCCACAUCUU | 21 | UAUCUCUGUGUCG | 23 | UGGUUGACCAUAGAACAUGCGC | 27 | UAUGUAAUAUGGUCCACAUCUU | 14 |
| hsa-mir-381 | UACUUAAAGCGAGGUUGCCCUUUGUAUAUUCGGUUUAUUGACAUGGAAUAUACAAGGGCAAGCUCUCUGUGAGUA | 24 | UCGGUUUAUUGACAUGGAA | 26 | AGCGAGGUUGCCCUUUGUAUAU | 27 | UAUACAAGGGCAAGCUCUCUGU | 23 |
| hsa-mir-382 | UACUUGAAGAGAAGUUGUUCGUGGUGGAUUCGCUUUACUUAUGACGAAUCAUUCACGGACAACACUUUUUUCAGUA | 21 | CUUUACUUAUGACG | 14 | GAAGUUGUUCGUGGUGGAUUCG | 41 | AAUCAUUCACGGACAACACUU | 10 |
| hsa-mir-383 | CUCCUCAGAUCAGAAGGUGAUUGUGGCUUUGGGUGGAUAUUAAUCAGCCACAGCACUGCCUGGUCAGAAAGAG | 29 | UUGGGUGGAUAUUAAUCAGCC | 29 | AGAUCAGAAGGUGAUUGUGGCU | 36 | ACAGCACUGCCUGGUCAGA | 26 |
| hsa-mir-409 | UGGUACUCGGGGAGAGGUUACCCGAGCAACUUUGCAUCUGGACGACGAAUGUUGCUCGGUGAACCCCUUUUCGGUAUCA | 29 | CUGGACGAC | 33 | AGGUUACCCGAGCAACUUUGCAU | 22 | GAAUGUUGCUCGGUGAACCCCU | 27 |
| hsa-mir-410 | GGUACCUGAGAAGAGGUUGUCUGUGAUGAGUUCGCUUUUAUUAAUGACGAAUAUAACACAGAUGGCCUGUUUUCAGUACC | 25 | CUUUUAUUAAUGACG | 13 | AGGUUGUCUGUGAUGAGUUCG | 38 | AAUAUAACACAGAUGGCCUGU | 19 |
| hsa-mir-411 | UGGUACUUGGAGAGAUAGUAGACCGUAUAGCGUACGCUUUAUCUGUGACGUAUGUAACACGGUCCACUAACCCUCAGUAUCAAAUCCAUCCCCGAG | 22 | CUUUAUCUGUGACG | 21 | UAGUAGACCGUAUAGCGUACG | 29 | UAUGUAACACGGUCCACUAACC | 14 |
| hsa-mir-412 | CUGGGGUACGGGGAUGGAUGGUCGACCAGUUGGAAAGUAAUUGUUUCUAAUGUACUUCACCUGGUCCACUAGCCGUCCGUAUCCGCUGCAG | 30 | UGUUUCUAAUGU | 17 | UGGUCGACCAGUUGGAAAGUAAU | 30 | ACUUCACCUGGUCCACUAGCCGU | 17 |
| hsa-mir-423 | AUAAAGGAAGUUAGGCUGAGGGGCAGAGAGCGAGACUUUUCUAUUUUCCAAAAGCUCGGUCUGAGGCCCCUCAGUCUUGCUUCCUAACCCGCGC | 27 | UCUAUUUUCCAAA | 0 | UGAGGGGCAGAGAGCGAGACUUU | 43 | AGCUCGGUCUGAGGCCCCUCAGU | 30 |
| hsa-mir-424 | CGAGGGGAUACAGCAGCAAUUCAUGUUUUGAAGUGUUCUAAAUGGUUCAAAACGUGAGGCGCUGCUAUACCCCCUCGUGGGGAAGGUAGAAGGUGGGG | 34 | GUGUUCUAAAUGGUU | 27 | CAGCAGCAAUUCAUGUUUUGAA | 18 | CAAAACGUGAGGCGCUGCUAU | 29 |
| hsa-mir-425 | GAAAGCGCUUUGGAAUGACACGAUCACUCCCGUUGAGUGGGCACCCGAGAAGCCAUCGGGAAUGUCGUGUCCGCCCAGUGCUCUUUC | 29 | GUGGGCACCCGAGAAGCC | 39 | AAUGACACGAUCACUCCCGUUGA | 17 | AUCGGGAAUGUCGUGUCCGCCC | 32 |
| hsa-mir-431 | UCCUGCUUGUCCUGCGAGGUGUCUUGCAGGCCGUCAUGCAGGCCACACUGACGGUAACGUUGCAGGUCGUCUUGCAGGGCUUCUCGCAAGACGACAUCCUCAUCACCAACGACG | 27 | GGCCACACUGACGGUAACGUUG | 32 | UGUCUUGCAGGCCGUCAUGCA | 29 | CAGGUCGUCUUGCAGGGCUUCU | 32 |
| hsa-mir-432 | UGACUCCUCCAGGUCUUGGAGUAGGUCAUUGGGUGGAUCCUCUAUUUCCUUACGUGGGCCACUGGAUGGCUCCUCCAUGUCUUGGAGUAGAUCA | 28 | AUCCUCUAUUUCCUUACGUGGGCCA | 16 | UCUUGGAGUAGGUCAUUGGGUGG | 43 | CUGGAUGGCUCCUCCAUGUCU | 24 |
| hsa-mir-433 | CCGGGGAGAAGUACGGUGAGCCUGUCAUUAUUCAGAGAGGCUAGAUCCUCUGUGUUGAGAAGGAUCAUGAUGGGCUCCUCGGUGUUCUCCAGG | 33 | AGAGAGGCUAGAUCCUCUGUGUUGAGAAGG | 37 | UACGGUGAGCCUGUCAUUAUUC | 23 | AUCAUGAUGGGCUCCUCGGUGU | 32 |
| hsa-mir-449b | UGACCUGAAUCAGGUAGGCAGUGUAUUGUUAGCUGGCUGCUUGGGUCAAGUCAGCAGCCACAACUACCCUGCCACUUGCUUCUGGAUAAAUUCUUCU | 24 | UGCUUGGGUCAAGUCAG | 35 | AGGCAGUGUAUUGUUAGCUGGC | 36 | CAGCCACAACUACCCUGCCACU | 9 |
| hsa-mir-449c | GCUGGGAUGUGUCAGGUAGGCAGUGUAUUGCUAGCGGCUGUUAAUGAUUUUAACAGUUGCUAGUUGCACUCCUCUCUGUUGCAUUCAGAAGC | 28 | UAAUGAUUUUAACAG | 13 | UAGGCAGUGUAUUGCUAGCGGCUGU | 36 | UUGCUAGUUGCACUCCUCUCUGU | 17 |
| hsa-mir-450a-1 | AAACGAUACUAAACUGUUUUUGCGAUGUGUUCCUAAUAUGCACUAUAAAUAUAUUGGGAACAUUUUGCAUGUAUAGUUUUGUAUCAAUAUA | 15 | GCACUAUAAAUAU | 8 | UUUUGCGAUGUGUUCCUAAUAU | 18 | AUUGGGAACAUUUUGCAUGUAU | 23 |
| hsa-mir-450a-2 | CCAAAGAAAGAUGCUAAACUAUUUUUGCGAUGUGUUCCUAAUAUGUAAUAUAAAUGUAUUGGGGACAUUUUGCAUUCAUAGUUUUGUAUCAAUAAUAUGG | 18 | GUAAUAUAAAUGU | 15 | UUUUGCGAUGUGUUCCUAAUAU | 18 | AUUGGGGACAUUUUGCAUUCAU | 23 |
| hsa-mir-450b | GCAGAAUUAUUUUUGCAAUAUGUUCCUGAAUAUGUAAUAUAAGUGUAUUGGGAUCAUUUUGCAUCCAUAGUUUUGUAU | 18 | UGUAAUAUAAGUGUA | 20 | UUUUGCAAUAUGUUCCUGAAUA | 14 | UUGGGAUCAUUUUGCAUCCAUA | 18 |
| hsa-mir-452 | GCUAAGCACUUACAACUGUUUGCAGAGGAAACUGAGACUUUGUAACUAUGUCUCAGUCUCAUCUGCAAAGAAGUAAGUGCUUUGC | 21 | GACUUUGUAACUAUGUCUCAGU | 18 | AACUGUUUGCAGAGGAAACUGA | 27 | CUCAUCUGCAAAGAAGUAAGUG | 23 |
| hsa-mir-454 | UCUGUUUAUCACCAGAUCCUAGAACCCUAUCAAUAUUGUCUCUGCUGUGUAAAUAGUUCUGAGUAGUGCAAUAUUGCUUAUAGGGUUUUGGUGUUUGGAAAGAACAAUGGGCAGG | 23 | UGUGUAAAUAGUUCUGAG | 28 | ACCCUAUCAAUAUUGUCUCUGC | 9 | UAGUGCAAUAUUGCUUAUAGGGU | 26 |
| hsa-mir-455 | UCCCUGGCGUGAGGGUAUGUGCCUUUGGACUACAUCGUGGAAGCCAGCACCAUGCAGUCCAUGGGCAUAUACACUUGCCUCAAGGCCUAUGUCAUC | 26 | UGGAAGCCAGCACCAU | 25 | UAUGUGCCUUUGGACUACAUCG | 23 | GCAGUCCAUGGGCAUAUACAC | 24 |
| hsa-mir-483 | GAGGGGGAAGACGGGAGGAAAGAAGGGAGUGGUUCCAUCACGCCUCCUCACUCCUCUCCUCCCGUCUUCUCCUCUC | 28 | UGGUUCCAUCACGCCUCC | 17 | AAGACGGGAGGAAAGAAGGGAG | 50 | UCACUCCUCUCCUCCCGUCUU | 5 |
| hsa-mir-485 | ACUUGGAGAGAGGCUGGCCGUGAUGAAUUCGAUUCAUCAAAGCGAGUCAUACACGGCUCUCCUCUCUUUUAGU | 25 | GAUUCAUCAAAGCGA | 20 | AGAGGCUGGCCGUGAUGAAUUC | 36 | GUCAUACACGGCUCUCCUCUCU | 14 |
| hsa-mir-486-1 | GCAUCCUGUACUGAGCUGCCCCGAGGCCCUUCAUGCUGCCCAGCUCGGGGCAGCUCAGUACAGGAUAC | 28 | GCCCUUCAUGCUGCCCAGCU | 20 | UCCUGUACUGAGCUGCCCCGAG | 27 | CGGGGCAGCUCAGUACAGGAU | 38 |
| hsa-mir-486-2 | UCCUGUACUGAGCUGCCCCGAGCUGGGCAGCAUGAAGGGCCUCGGGGCAGCUCAGUACAGGAUG | 36 | CUGGGCAGCAUGAAGGGCCU | 40 | UCCUGUACUGAGCUGCCCCGAG | 27 | CGGGGCAGCUCAGUACAGGAU | 38 |
| hsa-mir-487a | GGUACUUGAAGAGUGGUUAUCCCUGCUGUGUUCGCUUAAUUUAUGACGAAUCAUACAGGGACAUCCAGUUUUUCAGUAUC | 23 | CUUAAUUUAUGACG | 14 | GUGGUUAUCCCUGCUGUGUUCG | 32 | AAUCAUACAGGGACAUCCAGUU | 18 |
| hsa-mir-487b | UUGGUACUUGGAGAGUGGUUAUCCCUGUCCUGUUCGUUUUGCUCAUGUCGAAUCGUACAGGGUCAUCCACUUUUUCAGUAUCAA | 23 | UUUUGCUCAUGUCG | 21 | GUGGUUAUCCCUGUCCUGUUCG | 27 | AAUCGUACAGGGUCAUCCACUU | 18 |
| hsa-mir-488 | GAGAAUCAUCUCUCCCAGAUAAUGGCACUCUCAAACAAGUUUCCAAAUUGUUUGAAAGGCUAUUUCUUGGUCAGAUGACUCUC | 17 | ACAAGUUUCCAAAUUGU | 12 | CCCAGAUAAUGGCACUCUCAA | 14 | UUGAAAGGCUAUUUCUUGGUC | 24 |
| hsa-mir-489 | GUGGCAGCUUGGUGGUCGUAUGUGUGACGCCAUUUACUUGAACCUUUAGGAGUGACAUCACAUAUACGGCAGCUAAACUGCUAC | 26 | ACUUGAACCUUUAGGA | 19 | GGUCGUAUGUGUGACGCCAUUU | 32 | GUGACAUCACAUAUACGGCAGC | 23 |
| hsa-mir-490 | UGGAGGCCUUGCUGGUUUGGAAAGUUCAUUGUUCGACACCAUGGAUCUCCAGGUGGGUCAAGUUUAGAGAUGCACCAACCUGGAGGACUCCAUGCUGUUGAGCUGUUCACAAGCAGCGGACACUUCCA | 28 | CAAGUUUAGAGAUGCAC | 24 | CCAUGGAUCUCCAGGUGGGU | 35 | CAACCUGGAGGACUCCAUGCUG | 27 |
| hsa-mir-491 | UUGACUUAGCUGGGUAGUGGGGAACCCUUCCAUGAGGAGUAGAACACUCCUUAUGCAAGAUUCCCUUCUACCUGGCUGGGUUGG | 29 | AGUAGAACACUC | 17 | AGUGGGGAACCCUUCCAUGAGG | 36 | CUUAUGCAAGAUUCCCUUCUAC | 9 |
| hsa-mir-493 | CUGGCCUCCAGGGCUUUGUACAUGGUAGGCUUUCAUUCAUUCGUUUGCACAUUCGGUGAAGGUCUACUGUGUGCCAGGCCCUGUGCCAG | 28 | CAUUCGUUUGCACAUUCGG | 21 | UUGUACAUGGUAGGCUUUCAUU | 23 | UGAAGGUCUACUGUGUGCCAGG | 36 |
| hsa-mir-494 | GAUACUCGAAGGAGAGGUUGUCCGUGUUGUCUUCUCUUUAUUUAUGAUGAAACAUACACGGGAAACCUCUUUUUUAGUAUC | 21 | UUAUUUAUGA | 10 | AGGUUGUCCGUGUUGUCUUCUCU | 26 | UGAAACAUACACGGGAAACCUC | 18 |
| hsa-mir-495 | UGGUACCUGAAAAGAAGUUGCCCAUGUUAUUUUCGCUUUAUAUGUGACGAAACAAACAUGGUGCACUUCUUUUUCGGUAUCA | 20 | CUUUAUAUGUGACG | 21 | GAAGUUGCCCAUGUUAUUUUCG | 23 | AAACAAACAUGGUGCACUUCUU | 14 |
| hsa-mir-497 | CCACCCCGGUCCUGCUCCCGCCCCAGCAGCACACUGUGGUUUGUACGGCACUGUGGCCACGUCCAAACCACACUGUGGUGUUAGAGCGAGGGUGGGGGAGGCACCGCCGAGG | 33 | ACGGCACUGUGGCCACGUC | 32 | CAGCAGCACACUGUGGUUUGU | 29 | CAAACCACACUGUGGUGUUAGA | 23 |
| hsa-mir-499a | GCCCUGUCCCCUGUGCCUUGGGCGGGCGGCUGUUAAGACUUGCAGUGAUGUUUAACUCCUCUCCACGUGAACAUCACAGCAAGUCUGUGCUGCUUCCCGUCCCUACGCUGCCUGGGCAGGGU | 28 | AACUCCUCUCCACGUG | 13 | UUAAGACUUGCAGUGAUGUUU | 24 | AACAUCACAGCAAGUCUGUGCU | 18 |
| hsa-mir-499b | GGAAGCAGCACAGACUUGCUGUGAUGUUCACGUGGAGAGGAGUUAAACAUCACUGCAAGUCUUAACAGCCGCC | 27 | CGUGGAGAGGAGUUA | 47 | ACAGACUUGCUGUGAUGUUCA | 24 | AACAUCACUGCAAGUCUUAACA | 9 |
| hsa-mir-500a | GCUCCCCCUCUCUAAUCCUUGCUACCUGGGUGAGAGUGCUGUCUGAAUGCAAUGCACCUGGGCAAGGAUUCUGAGAGCGAGAGC | 29 | GUGCUGUCUGAAUGCA | 31 | UAAUCCUUGCUACCUGGGUGAGA | 26 | AUGCACCUGGGCAAGGAUUCUG | 32 |
| hsa-mir-500b | CCCCCUCUCUAAUCCUUGCUACCUGGGUGAGAGUGCUUUCUGAAUGCAGUGCACCCAGGCAAGGAUUCUGCAAGGGGGA | 28 | GAGAGUGCUUUCUGAAUGCAGU | 32 | AAUCCUUGCUACCUGGGU | 22 | GCACCCAGGCAAGGAUUCUG | 30 |
| hsa-mir-7-1 | UUGGAUGUUGGCCUAGUUCUGUGUGGAAGACUAGUGAUUUUGUUGUUUUUAGAUAACUAAAUCGACAACAAAUCACAGUCUGCCAUAUGGCACAGGCCAUGCCUCUACAG | 23 | UUUUAGAUAACUAAAUCGA | 11 | UGGAAGACUAGUGAUUUUGUUGU | 30 | CAACAAAUCACAGUCUGCCAUA | 9 |
| hsa-mir-7-2 | CUGGAUACAGAGUGGACCGGCUGGCCCCAUCUGGAAGACUAGUGAUUUUGUUGUUGUCUUACUGCGCUCAACAACAAAUCCCAGUCUACCUAAUGGUGCCAGCCAUCGCA | 24 | UGUCUUACUGCGCUCAA | 18 | UGGAAGACUAGUGAUUUUGUUGU | 30 | CAACAAAUCCCAGUCUACCUAA | 5 |
| hsa-mir-9-1 | CGGGGUUGGUUGUUAUCUUUGGUUAUCUAGCUGUAUGAGUGGUGUGGAGUCUUCAUAAAGCUAGAUAACCGAAAGUAAAAAUAACCCCA | 26 | GUGGUGUGGAGUCUUC | 44 | UCUUUGGUUAUCUAGCUGUAUGA | 22 | AUAAAGCUAGAUAACCGAAAGU | 18 |
| hsa-mir-9-2 | GGAAGCGAGUUGUUAUCUUUGGUUAUCUAGCUGUAUGAGUGUAUUGGUCUUCAUAAAGCUAGAUAACCGAAAGUAAAAACUCCUUCA | 22 | GUGUAUUGGUCUUC | 29 | UCUUUGGUUAUCUAGCUGUAUGA | 22 | AUAAAGCUAGAUAACCGAAAGU | 18 |
| hsa-mir-92a-1 | CUUUCUACACAGGUUGGGAUCGGUUGCAAUGCUGUGUUUCUGUAUGGUAUUGCACUUGUCCCGGCCUGUUGAGUUUGG | 29 | GUGUUUCUGUAUGG | 36 | AGGUUGGGAUCGGUUGCAAUGCU | 39 | UAUUGCACUUGUCCCGGCCUGU | 23 |
| hsa-mir-92a-2 | UCAUCCCUGGGUGGGGAUUUGUUGCAUUACUUGUGUUCUAUAUAAAGUAUUGCACUUGUCCCGGCCUGUGGAAGA | 27 | UUGUGUUCUAUAUAAAG | 18 | GGGUGGGGAUUUGUUGCAUUAC | 41 | UAUUGCACUUGUCCCGGCCUGU | 23 |
| hsa-mir-92b | CGGGCCCCGGGCGGGCGGGAGGGACGGGACGCGGUGCAGUGUUGUUUUUUCCCCCGCCAAUAUUGCACUCGUCCCGGCCUCCGGCCCCCCCGGCCC | 35 | UUGUUUUUUCCCCCGCCAA | 11 | AGGGACGGGACGCGGUGCAGUG | 55 | UAUUGCACUCGUCCCGGCCUCC | 18 |
| hsa-mir-93 | CUGGGGGCUCCAAAGUGCUGUUCGUGCAGGUAGUGUGAUUACCCAACCUACUGCUGAGCUAGCACUUCCCGAGCCCCCGG | 29 | UGUGAUUACCCAACCU | 13 | CAAAGUGCUGUUCGUGCAGGUAG | 35 | ACUGCUGAGCUAGCACUUCCCG | 23 |
| hsa-mir-9-3 | GGAGGCCCGUUUCUCUCUUUGGUUAUCUAGCUGUAUGAGUGCCACAGAGCCGUCAUAAAGCUAGAUAACCGAAAGUAGAAAUGAUUCUCA | 23 | GUGCCACAGAGCCGUC | 31 | UCUUUGGUUAUCUAGCUGUAUGA | 22 | AUAAAGCUAGAUAACCGAAAGU | 18 |
| hsa-mir-95 | AACACAGUGGGCACUCAAUAAAUGUCUGUUGAAUUGAAAUGCGUUACAUUCAACGGGUAUUUAUUGAGCACCCACUCUGUG | 21 | GAAAUGCGUUACA | 23 | UCAAUAAAUGUCUGUUGAAUU | 14 | UUCAACGGGUAUUUAUUGAGCA | 23 |
| hsa-mir-96 | UGGCCGAUUUUGGCACUAGCACAUUUUUGCUUGUGUCUCUCCGCUCUGAGCAAUCAUGUGCAGUGCCAAUAUGGGAAA | 24 | UGUGUCUCUCCGCUCUGAGC | 25 | UUUGGCACUAGCACAUUUUUGCU | 17 | AAUCAUGUGCAGUGCCAAUAUG | 23 |
| hsa-mir-98 | AGGAUUCUGCUCAUGCCAGGGUGAGGUAGUAAGUUGUAUUGUUGUGGGGUAGGGAUAUUAGGCCCCAAUUAGAAGAUAACUAUACAACUUACUACUUUCCCUGGUGUGUGGCAUAUUCA | 27 | GUGGGGUAGGGAUAUUAGGCCCCAAUUAGAAGAUAA | 33 | UGAGGUAGUAAGUUGUAUUGUU | 32 | CUAUACAACUUACUACUUUCCC | 0 |
| hsa-mir-99a | CCCAUUGGCAUAAACCCGUAGAUCCGAUCUUGUGGUGAAGUGGACCGCACAAGCUCGCUUCUAUGGGUCUGUGUCAGUGUG | 28 | GUGAAGUGGACCGCA | 40 | AACCCGUAGAUCCGAUCUUGUG | 23 | CAAGCUCGCUUCUAUGGGUCUG | 27 |
| hsa-mir-99b | GGCACCCACCCGUAGAACCGACCUUGCGGGGCCUUCGCCGCACACAAGCUCGUGUCUGUGGGUCCGUGUC | 30 | GGGCCUUCGCCGCACA | 31 | CACCCGUAGAACCGACCUUGCG | 23 | CAAGCUCGUGUCUGUGGGUCCG | 36 |
