## Supplementary material for "Guanine content of microRNAs is associated with their tumor suppressive and oncogenic roles in lung and breast cancers": Supp Table 2

Supplementary Table 2. Tumor suppressive miRs, oncomiRs, and the references from PubMed Database describing their function in breast and lung cancers. When search in PubMed had no results it appears as unknown function.

| **miRNA** | **Lung cancer** | **References** |  |  | **Breast cancer** | **References** |  |  |
| --- | --- | --- | --- | --- | --- | --- | --- | --- |
| **OncomiRs** | | | | | | | | |
| hsa-mir-328 | **oncomiR** | <https://www.ncbi.nlm.nih.gov/pubmed/21448905> | | | **unknown** |  |  |  |
| hsa-mir-197 | **oncomiR** | <https://www.ncbi.nlm.nih.gov/pubmed/24488097> | | | **unknown** |  |  |  |
| hsa-mir-92b | **oncomiR** | <https://www.ncbi.nlm.nih.gov/pubmed/24162673>  <https://www.ncbi.nlm.nih.gov/pubmed/24099768> | | | **unknown** |  |  |  |
| hsa-mir-483 | **oncomiR** | <https://www.ncbi.nlm.nih.gov/pubmed/24710410> | | | **unknown** |  |  |  |
| hsa-mir-23a | **oncomiR** | <https://www.ncbi.nlm.nih.gov/pubmed/24898878> | | | **oncomiR** | <https://www.ncbi.nlm.nih.gov/pubmed/29050223>  <https://www.ncbi.nlm.nih.gov/pubmed/23649631> | | |
| hsa-mir-25 | **oncomiR** | <https://www.ncbi.nlm.nih.gov/pubmed/26464659>  <https://www.ncbi.nlm.nih.gov/pubmed/25432132>  <https://www.ncbi.nlm.nih.gov/pubmed/25998847h> | | | **oncomiR** | <https://www.ncbi.nlm.nih.gov/pubmed/29310680> | | |
| hsa-mir-27a | **oncomiR** | <https://www.ncbi.nlm.nih.gov/pubmed/28370334>  <https://www.ncbi.nlm.nih.gov/pubmed/21460851> | | | **oncomiR** | <https://www.ncbi.nlm.nih.gov/pubmed/18006846/>  <https://www.ncbi.nlm.nih.gov/pubmed/23240057> | | |
| hsa-mir-196a | **oncomiR** | <https://www.ncbi.nlm.nih.gov/pubmed/22876840>  <https://www.ncbi.nlm.nih.gov/pubmed/27880728> | | | **oncomiR** | <https://www.ncbi.nlm.nih.gov/pubmed/26062455>  <https://www.ncbi.nlm.nih.gov/pubmed/27105503> | | |
| hsa-mir-93 | **oncomiR** | <https://www.ncbi.nlm.nih.gov/pubmed/29309884>  <https://www.ncbi.nlm.nih.gov/pubmed/28382150>  <https://www.ncbi.nlm.nih.gov/pubmed/24037530>  <https://www.ncbi.nlm.nih.gov/pubmed/23111389> | | | **oncomiR** | <https://www.ncbi.nlm.nih.gov/pubmed/28518139>  <https://www.ncbi.nlm.nih.gov/pubmed/25238878>  <https://www.ncbi.nlm.nih.gov/pubmed/23492819> | **Tumor suppressor miR** | <https://www.ncbi.nlm.nih.gov/pubmed/27840899> |
| hsa-mir-487a | **oncomiR** | <https://www.ncbi.nlm.nih.gov/pubmed/29330288> | | | **oncomiR** | <https://www.ncbi.nlm.nih.gov/pubmed/27019625> | | |
| hsa-mir-20a | **oncomiR** | <https://www.ncbi.nlm.nih.gov/pubmed/29375712>  <https://www.ncbi.nlm.nih.gov/pubmed/17384677> | | | **unknown** |  |  |  |
| hsa-mir-155 | **oncomiR** | <https://www.ncbi.nlm.nih.gov/pubmed/26548866> | | | **oncomiR** | <https://www.ncbi.nlm.nih.gov/pubmed/24152184> | | |
| hsa-mir-24 | **oncomiR** | <https://www.ncbi.nlm.nih.gov/pubmed/29231262>  <https://www.ncbi.nlm.nih.gov/pubmed/25725584> | | | **oncomiR** | <https://www.ncbi.nlm.nih.gov/pubmed/23418360>  <https://www.ncbi.nlm.nih.gov/pubmed/28490335>  <https://www.ncbi.nlm.nih.gov/pubmed/26044523> | | |
| hsa-mir-135b | **oncomiR** | <https://www.ncbi.nlm.nih.gov/pubmed/23695671> | | | **oncomiR** | <https://www.ncbi.nlm.nih.gov/pubmed/26934863> | | |
| hsa-mir-150 | **oncomiR** | <https://www.ncbi.nlm.nih.gov/pubmed/28891208>  <https://www.ncbi.nlm.nih.gov/pubmed/27976702>  <https://www.ncbi.nlm.nih.gov/pubmed/23747308>  <https://www.ncbi.nlm.nih.gov/pubmed/24456795> | **Tumor suppressor miR** | <https://www.ncbi.nlm.nih.gov/pubmed/29286099>  <https://www.ncbi.nlm.nih.gov/pubmed/28108217> | **oncomiR** | <https://www.ncbi.nlm.nih.gov/pubmed/24312495> | | |
| hsa-mir-301a | **oncomiR** | <https://www.ncbi.nlm.nih.gov/pubmed/29113240> | | | **oncomiR** | <https://www.ncbi.nlm.nih.gov/pubmed/24315818>  <https://www.ncbi.nlm.nih.gov/pubmed/25311065> | | |
| hsa-mir-367 | **oncomiR** | <https://www.ncbi.nlm.nih.gov/pubmed/28656290>  <https://www.ncbi.nlm.nih.gov/pubmed/28000899> | | | **unknown** |  |  |  |
| hsa-mir-95 | **oncomiR** | <https://www.ncbi.nlm.nih.gov/pubmed/24835695> | | | **unknown** |  |  |  |
| hsa-mir-454 | **oncomiR** | <https://www.ncbi.nlm.nih.gov/pubmed/27261580> | | | **oncomiR** | <https://www.ncbi.nlm.nih.gov/pubmed/28795052> | | |
| hsa-mir-106a | **oncomiR** | <https://www.ncbi.nlm.nih.gov/pubmed/26097565> | | | **unknown** |  |  |  |
| hsa-mir-151a | **oncomiR** | <https://www.ncbi.nlm.nih.gov/pubmed/28759022> | | | **unknown** |  |  |  |
| hsa-mir-301b | **oncomiR** | <https://www.ncbi.nlm.nih.gov/pubmed/27352910> | | | **unknown** |  |  |  |
| hsa-mir-181c | **unknown** |  |  |  | **oncomiR** | <https://www.ncbi.nlm.nih.gov/pubmed/25695913> | | |
| hsa-mir-425 | **unknown** |  |  |  | **oncomiR** | <https://www.ncbi.nlm.nih.gov/pubmed/28038450> | | |
| hsa-mir-372 | **oncomiR** | <https://www.ncbi.nlm.nih.gov/pubmed/28440022> | | | **unknown** |  | | |
| **Tumor suppressor miRs** | | | | | | | | |
| hsa-mir-132 | **tumor suppressor miR** | <https://www.ncbi.nlm.nih.gov/pubmed/26543603>  <https://www.ncbi.nlm.nih.gov/pubmed/24626466 https://www.ncbi.nlm.nih.gov/pubmed/27735039> | | | **tumor suppressor miR** | <https://www.ncbi.nlm.nih.gov/pubmed/25450365>  <https://www.ncbi.nlm.nih.gov/pubmed/23399321>  <https://www.ncbi.nlm.nih.gov/pubmed/28816236> | | |
| hsa-mir-33b | **tumor suppressor miR** | <https://www.ncbi.nlm.nih.gov/pubmed/26459797> | | | **tumor suppressor miR** | <https://www.ncbi.nlm.nih.gov/pubmed/25919570> | | |
| hsa-mir-139 | **tumor suppressor miR** | <https://www.ncbi.nlm.nih.gov/pubmed/26256448>  <https://www.ncbi.nlm.nih.gov/pubmed/26497851> | | | **tumor suppressor miR** | <https://www.ncbi.nlm.nih.gov/pubmed/27864119>  <https://www.ncbi.nlm.nih.gov/pubmed/26299922>  <https://www.ncbi.nlm.nih.gov/pubmed/26079880>  <https://www.ncbi.nlm.nih.gov/pubmed/24158791> | | |
| hsa-mir-149 | **tumor suppressor miR** | <https://www.ncbi.nlm.nih.gov/pubmed/29130108>  <https://www.ncbi.nlm.nih.gov/pubmed/23762558> | | | **tumor suppressor miR** | <https://www.ncbi.nlm.nih.gov/pubmed/29270025>  <https://www.ncbi.nlm.nih.gov/pubmed/2460843> | | |
| hsa-mir-193a | **tumor suppressor miR** | <https://www.ncbi.nlm.nih.gov/pubmed/29183007>  <https://www.ncbi.nlm.nih.gov/pubmed/25391651> | | | **tumor suppressor miR** | <https://www.ncbi.nlm.nih.gov/pubmed/29016617> | | |
| hsa-mir-200c | **tumor suppressor miR** | <https://www.ncbi.nlm.nih.gov/pubmed/29321091>  <https://www.ncbi.nlm.nih.gov/pubmed/28727734> | | | **tumor suppressor miR** | <https://www.ncbi.nlm.nih.gov/pubmed/29162923>  <https://www.ncbi.nlm.nih.gov/pubmed/29155146>  <https://www.ncbi.nlm.nih.gov/pubmed/28829888> | | |
| hsa-mir-152 | **tumor suppressor miR** | <https://www.ncbi.nlm.nih.gov/pubmed/26823738>  <https://www.ncbi.nlm.nih.gov/pubmed/25190353>  <https://www.ncbi.nlm.nih.gov/pubmed/24780186>  <https://www.ncbi.nlm.nih.gov/pubmed/20696752>  <https://www.ncbi.nlm.nih.gov/pubmed/24997798> | | | **tumor suppressor miR** | <https://www.ncbi.nlm.nih.gov/pubmed/28653610>  <https://www.ncbi.nlm.nih.gov/pubmed/28247844>  <https://www.ncbi.nlm.nih.gov/pubmed/26392416>  <https://www.ncbi.nlm.nih.gov/pubmed/24368337>  <https://www.ncbi.nlm.nih.gov/pubmed/24710933> | | |
| hsa-mir-30c | **tumor suppressor miR** | <https://www.ncbi.nlm.nih.gov/pubmed/25249344>  <https://www.ncbi.nlm.nih.gov/pubmed/23988701>  <https://www.ncbi.nlm.nih.gov/pubmed/25119247> | | | **tumor suppressor miR** | <https://www.ncbi.nlm.nih.gov/pubmed/22701724> | | |
| hsa-mir-128 | **tumor suppressor miR** | <https://www.ncbi.nlm.nih.gov/pubmed/25001183> | | | **unknown** |  |  |  |
| hsa-mir-218 | **tumor suppressor miR** | <https://www.ncbi.nlm.nih.gov/pubmed/28830450>  <https://www.ncbi.nlm.nih.gov/pubmed/28429357>  <https://www.ncbi.nlm.nih.gov/pubmed/28192397>  <https://www.ncbi.nlm.nih.gov/pubmed/27057632>  <https://www.ncbi.nlm.nih.gov/pubmed/27633630> | | | **tumor suppressor miR** | <https://www.ncbi.nlm.nih.gov/pubmed/27109339>  <https://www.ncbi.nlm.nih.gov/pubmed/29378184> | | |
| hsa-mir-142 | **tumor suppressor miR** | <https://www.ncbi.nlm.nih.gov/pubmed/29131028>  <https://www.ncbi.nlm.nih.gov/pubmed/26617792> | | | **unknown** |  |  |  |
| hsa-mir-143 | **tumor suppressor miR** | <https://www.ncbi.nlm.nih.gov/pubmed/27602093>  <https://www.ncbi.nlm.nih.gov/pubmed/25003638>  <https://www.ncbi.nlm.nih.gov/pubmed/23904792> | | | **tumor suppressor miR** | <https://www.ncbi.nlm.nih.gov/pubmed/29360495>  <https://www.ncbi.nlm.nih.gov/pubmed/28559978>  <https://www.ncbi.nlm.nih.gov/pubmed/26618772>  <https://www.ncbi.nlm.nih.gov/pubmed/25248370>  <https://www.ncbi.nlm.nih.gov/pubmed/28511343> | | |
| hsa-mir-195 | **tumor suppressor miR** | <https://www.ncbi.nlm.nih.gov/pubmed/25840419>  <https://www.ncbi.nlm.nih.gov/pubmed/24891187>  <https://www.ncbi.nlm.nih.gov/pubmed/24874051>  <https://www.ncbi.nlm.nih.gov/pubmed/24486218> | | | **tumor suppressor miR** | <https://www.ncbi.nlm.nih.gov/pubmed/27133044>  <https://www.ncbi.nlm.nih.gov/pubmed/26632252>  <https://www.ncbi.nlm.nih.gov/pubmed/24402230>  <https://www.ncbi.nlm.nih.gov/pubmed/21350001> | | |
| hsa-mir-486 | **tumor suppressor miR** | <https://www.ncbi.nlm.nih.gov/pubmed/23980150>  <https://www.ncbi.nlm.nih.gov/pubmed/23474761> | | | **tumor suppressor miR** | <https://www.ncbi.nlm.nih.gov/pubmed/25104088> | | |
| hsa-mir-30e | **tumor suppressor miR** | <https://www.ncbi.nlm.nih.gov/pubmed/29174979>  <https://www.ncbi.nlm.nih.gov/pubmed/27992364> | **oncomiR** | <https://www.ncbi.nlm.nih.gov/pubmed/28653805> | **tumor suppressor miR** | <https://www.ncbi.nlm.nih.gov/pubmed/29162879> | | |
| hsa-mir-99a | **tumor suppressor miR** | <https://www.ncbi.nlm.nih.gov/pubmed/26986073>  <https://www.ncbi.nlm.nih.gov/pubmed/25663868>  <https://www.ncbi.nlm.nih.gov/pubmed/25187230> | | | **tumor suppressor miR** | <https://www.ncbi.nlm.nih.gov/pubmed/27212167>  <https://www.ncbi.nlm.nih.gov/pubmed/26417931>  <https://www.ncbi.nlm.nih.gov/pubmed/24637915>  <https://www.ncbi.nlm.nih.gov/pubmed/25348507> | | |
| hsa-mir-342 | **tumor suppressor miR** | <https://www.ncbi.nlm.nih.gov/pubmed/29107102>  <https://www.ncbi.nlm.nih.gov/pubmed/27670444>  <https://www.ncbi.nlm.nih.gov/pubmed/25663460> | | | **tumor suppressor miR** | <https://www.ncbi.nlm.nih.gov/pubmed/26919240> | | |
| hsa-mir-30a | **tumor suppressor miR** | <https://www.ncbi.nlm.nih.gov/pubmed/28678320>  <https://www.ncbi.nlm.nih.gov/pubmed/26837415>  <https://www.ncbi.nlm.nih.gov/pubmed/26025408>  <https://www.ncbi.nlm.nih.gov/pubmed/21633953> | | | **tumor suppressor miR** | <https://www.ncbi.nlm.nih.gov/pubmed/23851509>  <https://www.ncbi.nlm.nih.gov/pubmed/28765900>  <https://www.ncbi.nlm.nih.gov/pubmed/27003255>  <https://www.ncbi.nlm.nih.gov/pubmed/22476851>  <https://www.ncbi.nlm.nih.gov/pubmed/24508260> | | |
| hsa-mir-200b | **tumor suppressor miR** | <https://www.ncbi.nlm.nih.gov/pubmed/27356635> | | | **tumor suppressor miR** | <https://www.ncbi.nlm.nih.gov/pubmed/29156719>  <https://www.ncbi.nlm.nih.gov/pubmed/28692034>  <https://www.ncbi.nlm.nih.gov/pubmed/27276064>  <https://www.ncbi.nlm.nih.gov/pubmed/24925028> | | |
| hsa-mir-125a | **tumor suppressor miR** | <https://www.ncbi.nlm.nih.gov/pubmed/28631574>  <https://www.ncbi.nlm.nih.gov/pubmed/25998575>  <https://www.ncbi.nlm.nih.gov/pubmed/24044511>  <https://www.ncbi.nlm.nih.gov/pubmed/19702827>  <https://www.ncbi.nlm.nih.gov/pubmed/21777146> | | | **tumor suppressor miR** | <https://www.ncbi.nlm.nih.gov/pubmed/19875930> | | |
| hsa-mir-26b | **tumor suppressor miR** | <https://www.ncbi.nlm.nih.gov/pubmed/27078844>  <https://www.ncbi.nlm.nih.gov/pubmed/26827826>  <https://www.ncbi.nlm.nih.gov/pubmed/26744864> | | | **tumor suppressor miR** | <https://www.ncbi.nlm.nih.gov/pubmed/24753748>  <https://www.ncbi.nlm.nih.gov/pubmed/23374284>  <https://www.ncbi.nlm.nih.gov/pubmed/21510944>  <https://www.ncbi.nlm.nih.gov/pubmed/23939832> | | |
| hsa-mir-34b | **tumor suppressor miR** | <https://www.ncbi.nlm.nih.gov/pubmed/22047961> | | | **tumor suppressor miR** | <https://www.ncbi.nlm.nih.gov/pubmed/22113133> | | |
| hsa-mir-146a | **tumor suppressor miR** | <https://www.ncbi.nlm.nih.gov/pubmed/27494902>  <https://www.ncbi.nlm.nih.gov/pubmed/23555954> | | | **tumor suppressor miR** | <https://www.ncbi.nlm.nih.gov/pubmed/27175941>  <https://www.ncbi.nlm.nih.gov/pubmed/18504431> | **oncomiR** | <https://www.ncbi.nlm.nih.gov/pubmed/25123132> |
| hsa-mir-212 | **tumor suppressor miR** | <https://www.ncbi.nlm.nih.gov/pubmed/28791372>  <https://www.ncbi.nlm.nih.gov/pubmed/25435090> | **oncomiR** | <https://www.ncbi.nlm.nih.gov/pubmed/22357618> | **tumor suppressor miR** | <https://www.ncbi.nlm.nih.gov/pubmed/29216628> | | |
| hsa-mir-204 | **tumor suppressor miR** | <https://www.ncbi.nlm.nih.gov/pubmed/27323056>  <https://www.ncbi.nlm.nih.gov/pubmed/25412236>  <https://www.ncbi.nlm.nih.gov/pubmed/25157435>  <https://www.ncbi.nlm.nih.gov/pubmed/28534958>  <https://www.ncbi.nlm.nih.gov/pubmed/26935060> | | | **tumor suppressor miR** | <https://www.ncbi.nlm.nih.gov/pubmed/28534958>  <https://www.ncbi.nlm.nih.gov/pubmed/26408179>  <https://www.ncbi.nlm.nih.gov/pubmed/26191195> | | |
| hsa-mir-34a | **tumor suppressor miR** | <https://www.ncbi.nlm.nih.gov/pubmed/25501507>  <https://www.ncbi.nlm.nih.gov/pubmed/26652031> | | | **tumor suppressor miR** | <https://www.ncbi.nlm.nih.gov/pubmed/22623155>  <https://www.ncbi.nlm.nih.gov/pubmed/28423566> | | |
| hsa-mir-219a | **unknown** |  |  |  | **tumor suppressor miR** | <https://www.ncbi.nlm.nih.gov/pubmed/29077787> | | |
| hsa-mir-340 | **tumor suppressor miR** | <https://www.ncbi.nlm.nih.gov/pubmed/25151966>  <https://www.ncbi.nlm.nih.gov/pubmed/27308439> | | | **tumor suppressor miR** | <https://www.ncbi.nlm.nih.gov/pubmed/28928895>  <https://www.ncbi.nlm.nih.gov/pubmed/28781664>  <https://www.ncbi.nlm.nih.gov/pubmed/26758430>  <https://www.ncbi.nlm.nih.gov/pubmed/21692045>  <https://www.ncbi.nlm.nih.gov/pubmed/26573744> | | |
| hsa-let-7e | **unknown** |  |  |  | **tumor suppressor miR** | <https://www.ncbi.nlm.nih.gov/pubmed/21969366> | | |
| hsa-let-7a | **tumor suppressor miR** | <https://www.ncbi.nlm.nih.gov/pubmed/23134218>  <https://www.ncbi.nlm.nih.gov/pubmed/20033209> | | | **tumor suppressor miR** | <https://www.ncbi.nlm.nih.gov/pubmed/29050238>  <https://www.ncbi.nlm.nih.gov/pubmed/26898455>  <https://www.ncbi.nlm.nih.gov/pubmed/25846193>  <https://www.ncbi.nlm.nih.gov/pubmed/22251626> | | |
| hsa-let-7d | **tumor suppressor miR** | <https://www.ncbi.nlm.nih.gov/pubmed/25477749/> | | | **tumor suppressor miR** | <https://www.ncbi.nlm.nih.gov/pubmed/25477749/> | | |
| hsa-let-7b | **tumor suppressor miR** | <https://www.ncbi.nlm.nih.gov/pubmed/22761738> | | | **tumor suppressor miR** | <https://www.ncbi.nlm.nih.gov/pubmed/28604753>  <https://www.ncbi.nlm.nih.gov/pubmed/25789066>  <https://www.ncbi.nlm.nih.gov/pubmed/23339187> | | |
| hsa-mir-30b | **tumor suppressor miR** | <https://www.ncbi.nlm.nih.gov/pubmed/26388700>  <https://www.ncbi.nlm.nih.gov/pubmed/25249344> | | | **tumor suppressor miR** | <https://www.ncbi.nlm.nih.gov/pubmed/22384020> | | |
| hsa-mir-202 | **tumor suppressor miR** | <https://www.ncbi.nlm.nih.gov/pubmed/28656198>  <https://www.ncbi.nlm.nih.gov/pubmed/27338052> | | | **unknown** |  |  |  |
| hsa-mir-383 | **tumor suppressor miR** | <https://www.ncbi.nlm.nih.gov/pubmed/28927114>  <https://www.ncbi.nlm.nih.gov/pubmed/27862077>  <https://www.ncbi.nlm.nih.gov/pubmed/27551765> | | | **unknown** |  |  |  |
| hsa-let-7c | **tumor suppressor miR** | <https://www.ncbi.nlm.nih.gov/pubmed/23981581>  <https://www.ncbi.nlm.nih.gov/pubmed/23850892>  <https://www.ncbi.nlm.nih.gov/pubmed/23534758> | | | **tumor suppressor miR** | <https://www.ncbi.nlm.nih.gov/pubmed/28604753>  <https://www.ncbi.nlm.nih.gov/pubmed/26987290>  <https://www.ncbi.nlm.nih.gov/pubmed/28731186> | | |
| hsa-mir-138 | **tumor suppressor miR** | <https://www.ncbi.nlm.nih.gov/pubmed/28653608>  <https://www.ncbi.nlm.nih.gov/pubmed/28498463>  <https://www.ncbi.nlm.nih.gov/pubmed/24405893>  <https://www.ncbi.nlm.nih.gov/pubmed/23343715>  <https://www.ncbi.nlm.nih.gov/pubmed/26283050>  <https://www.ncbi.nlm.nih.gov/pubmed/26201895>  <https://www.ncbi.nlm.nih.gov/pubmed/27665963>  <https://www.ncbi.nlm.nih.gov/pubmed/27223073> | | | **tumor suppressor miR** | <https://www.ncbi.nlm.nih.gov/pubmed/26796277>  <https://www.ncbi.nlm.nih.gov/pubmed/28893536> | | |
| hsa-mir-1 | **tumor suppressor miR** | <https://www.ncbi.nlm.nih.gov/pubmed/24486107>  <https://www.ncbi.nlm.nih.gov/pubmed/28853613>  <https://www.ncbi.nlm.nih.gov/pubmed/18818206> | | | **tumor suppressor miR** | <https://www.ncbi.nlm.nih.gov/pubmed/28159933>  <https://www.ncbi.nlm.nih.gov/pubmed/26497855>  <https://www.ncbi.nlm.nih.gov/pubmed/26275461> | | |
| hsa-mir-485 | **tumor suppressor miR** | <https://www.ncbi.nlm.nih.gov/pubmed/27262438> | | | **tumor suppressor miR** | <https://www.ncbi.nlm.nih.gov/pubmed/27010860>  <https://www.ncbi.nlm.nih.gov/pubmed/23886178> | | |
| hsa-mir-134 | **tumor suppressor miR** | <https://www.ncbi.nlm.nih.gov/pubmed/28075475>  <https://www.ncbi.nlm.nih.gov/pubmed/27241841>  <https://www.ncbi.nlm.nih.gov/pubmed/27166267>  <https://www.ncbi.nlm.nih.gov/pubmed/23010597> | | | **tumor suppressor miR** | <https://www.ncbi.nlm.nih.gov/pubmed/28454346> | | |
| hsa-mir-379 | **unknown** |  |  |  | **tumor suppressor miR** | <https://www.ncbi.nlm.nih.gov/pubmed/23874748> | | |
| hsa-mir-491 | **tumor suppressor miR** | <https://www.ncbi.nlm.nih.gov/pubmed/27158341> | | | **tumor suppressor miR** | <https://www.ncbi.nlm.nih.gov/pubmed/25725194> | | |
| hsa-mir-432 | **tumor suppressor miR** | <https://www.ncbi.nlm.nih.gov/pubmed/26942465> | | | **unknown** |  |  |  |
| hsa-mir-185 | **tumor suppressor miR** | <https://www.ncbi.nlm.nih.gov/pubmed/26617940>  <https://www.ncbi.nlm.nih.gov/pubmed/19688090> | | | **tumor suppressor miR** | <https://www.ncbi.nlm.nih.gov/pubmed/25371748>  <https://www.ncbi.nlm.nih.gov/pubmed/25319390>  <https://www.ncbi.nlm.nih.gov/pubmed/25448984>  <https://www.ncbi.nlm.nih.gov/pubmed/20603620> | | |
| hsa-mir-296 | **tumor suppressor miR** | <https://www.ncbi.nlm.nih.gov/pubmed/26549165> | | | **tumor suppressor miR** | <https://www.ncbi.nlm.nih.gov/pubmed/21643016>  <https://www.ncbi.nlm.nih.gov/pubmed/24527800> | | |
| hsa-mir-335 | **tumor suppressor miR** | <https://www.ncbi.nlm.nih.gov/pubmed/29161765>  <https://www.ncbi.nlm.nih.gov/pubmed/23740614> | | | **tumor suppressor miR** | <https://www.ncbi.nlm.nih.gov/pubmed/28795314>  <https://www.ncbi.nlm.nih.gov/pubmed/25492484>  <https://www.ncbi.nlm.nih.gov/pubmed/25323813>  <https://www.ncbi.nlm.nih.gov/pubmed/21618216>  <https://www.ncbi.nlm.nih.gov/pubmed/21289068> | | |
| hsa-mir-193b | **tumor suppressor miR** | <https://www.ncbi.nlm.nih.gov/pubmed/22491710> | | | **tumor suppressor miR** | <https://www.ncbi.nlm.nih.gov/pubmed/25550792> | | |
| hsa-mir-449c | **tumor suppressor miR** | <https://www.ncbi.nlm.nih.gov/pubmed/23507140> | | | **unknown** |  |  |  |
| hsa-mir-126 | **tumor suppressor miR** | <https://www.ncbi.nlm.nih.gov/pubmed/27236384>  <https://www.ncbi.nlm.nih.gov/pubmed/21439283>  <https://www.ncbi.nlm.nih.gov/pubmed/20034472>  <https://www.ncbi.nlm.nih.gov/pubmed/18602365> | | | **tumor suppressor miR** | <https://www.ncbi.nlm.nih.gov/pubmed/26261534>  <https://www.ncbi.nlm.nih.gov/pubmed/23396050> | | |
| hsa-mir-186 | **tumor suppressor miR** | <https://www.ncbi.nlm.nih.gov/pubmed/29125477>  <https://www.ncbi.nlm.nih.gov/pubmed/27498924>  <https://www.ncbi.nlm.nih.gov/pubmed/24894676> | | | **unknown** |  |  |  |
| hsa-mir-488 | **tumor suppressor miR** | <https://www.ncbi.nlm.nih.gov/pubmed/28074905> | | | **unknown** |  |  |  |
| hsa-mir-7 | **tumor suppressor miR** | <https://www.ncbi.nlm.nih.gov/pubmed/27764812>  <https://www.ncbi.nlm.nih.gov/pubmed/26135959>  <https://www.ncbi.nlm.nih.gov/pubmed/25334070>  <https://www.ncbi.nlm.nih.gov/pubmed/21750649>  <https://www.ncbi.nlm.nih.gov/pubmed/24281003> | **oncomiR** | <https://www.ncbi.nlm.nih.gov/pubmed/25181544>  <https://www.ncbi.nlm.nih.gov/pubmed/20978205> | **tumor suppressor miR** | <https://www.ncbi.nlm.nih.gov/pubmed/28571043>  <https://www.ncbi.nlm.nih.gov/pubmed/25511742>  <https://www.ncbi.nlm.nih.gov/pubmed/25532106>  <https://www.ncbi.nlm.nih.gov/pubmed/22876288> | | |
| hsa-mir-34c | **tumor suppressor miR** | <https://www.ncbi.nlm.nih.gov/pubmed/26261507> | | | **tumor suppressor miR** | <https://www.ncbi.nlm.nih.gov/pubmed/27698902> | | |
| hsa-mir-122 | **tumor suppressor miR** | <https://www.ncbi.nlm.nih.gov/pubmed/26604787> | | | **tumor suppressor miR** | <https://www.ncbi.nlm.nih.gov/pubmed/23056576> | | |
| hsa-mir-376a | **tumor suppressor miR** | <https://www.ncbi.nlm.nih.gov/pubmed/28741879> | | | **unknown** |  |  |  |
| hsa-mir-29a | **tumor suppressor miR** | <https://www.ncbi.nlm.nih.gov/pubmed/28521487>  <https://www.ncbi.nlm.nih.gov/pubmed/25171863> | | | **tumor suppressor miR** | <https://www.ncbi.nlm.nih.gov/pubmed/24289849> | **oncomiR** | <https://www.ncbi.nlm.nih.gov/pubmed/27555295> |
| hsa-mir-124 | **tumor suppressor miR** | <https://www.ncbi.nlm.nih.gov/pubmed/28927097>  <https://www.ncbi.nlm.nih.gov/pubmed/27376157>  <https://www.ncbi.nlm.nih.gov/pubmed/27251409>  <https://www.ncbi.nlm.nih.gov/pubmed/25531908>  <https://www.ncbi.nlm.nih.gov/pubmed/26935152>  <https://www.ncbi.nlm.nih.gov/pubmed/27073840> | | | **tumor suppressor miR** | <https://www.ncbi.nlm.nih.gov/pubmed/27842510>  <https://www.ncbi.nlm.nih.gov/pubmed/27748910>  <https://www.ncbi.nlm.nih.gov/pubmed/25731732>  <https://www.ncbi.nlm.nih.gov/pubmed/23816858>  <https://www.ncbi.nlm.nih.gov/pubmed/25085587> | | |
| hsa-mir-361 | **tumor suppressor miR** | <https://www.ncbi.nlm.nih.gov/pubmed/28837805>  <https://www.ncbi.nlm.nih.gov/pubmed/28485158>  <https://www.ncbi.nlm.nih.gov/pubmed/27779659>  <https://www.ncbi.nlm.nih.gov/pubmed/27164951> | | | **tumor suppressor miR** | <https://www.ncbi.nlm.nih.gov/pubmed/29132384> | | |
| hsa-mir-16 | **tumor suppressor miR** | <https://www.ncbi.nlm.nih.gov/pubmed/27712591>  <https://www.ncbi.nlm.nih.gov/pubmed/26064212>  <https://www.ncbi.nlm.nih.gov/pubmed/23954293> | | | **tumor suppressor miR** | <https://www.ncbi.nlm.nih.gov/pubmed/25261849> | | |
