## Supplementary material for "Guanine content of microRNAs is associated with their tumor suppressive and oncogenic roles in lung and breast cancers": Supp Table 3

Supplementary Table 3. G-enriched tumor suppressive miRs target genes. Shown are 30 most TL-G-enriched tumor suppressive miRs and their direct target genes in breast and lung cancers, as were described in the references from PubMed Database (Supplementary Table 2).

| **mir-132** | **mir-33b** | **mir-139** | **mir-149** | **mir-193a** | **mir-200c** | **mir-152** | **mir-30c** | **mir-218** | **mir-142** | **mir-128** | **mir-143** | **mir-195** | **mir-486** | **mir-99a** | **mir-30e** | **mir-342** | **mir-30a** | **mir-125a** | **mir-200b** | **mir-146a** | **mir-26b** | **mir-34b** | **mir-204** | **mir-212** | **mir-219a** | **mir-34a** | **mir-340** | **let-7e** | **let-7a** |
| --- | --- | --- | --- | --- | --- | --- | --- | --- | --- | --- | --- | --- | --- | --- | --- | --- | --- | --- | --- | --- | --- | --- | --- | --- | --- | --- | --- | --- | --- |
| **Lung cancer** | | | | | | | | | | | | | | | | | | | | | | | | | | | | | |
| SOX4 | ZEB1 | Met | FOXM1 | KRAS | LDHA | NRP1 | Rab18 | IL6 | PIK3CA | VEGFC | EGFR | CHEK1 | IGF1 | NOX4 | SIRT1 | AGR2 | Snai1 | STAT3 | FSCN1 | CCND1 | KPNA2 | Met | JAK2 | SOX4 |  | TGFBR2 | SKP2 |  | KRAS |
| ZEB2 |  |  |  | ERBB4 | HMGB1 | FGF2 | MTA1 | JAK3 | HMGB1 |  | Limk1 | HDGF | IGF1R | IGF1R |  | PTPRN | EYA2 | MTA1 |  | CCND2 | MIEN1 |  | NUAK1 |  |  | MDM4 | PUM1 |  | HMGA2 |
| Smad2 |  |  |  |  | USP25 | ADAM17 |  | CDCP1 |  |  | CD44v3 | IGF1R | PIK3R1 | AKT1 |  | RAP2B | IGF1R |  |  |  | PTGS2 |  | SIX1 |  |  | SERPINE1 | PUM2 |  | NIRF |
|  |  |  |  |  |  |  |  | Slug |  |  |  | MYB | ARHGAP5 |  |  |  |  |  |  |  |  |  | ATF2 |  |  | EGFR |  |  |  |
|  |  |  |  |  |  |  |  | ZEB2 |  |  |  |  |  |  |  |  |  |  |  |  |  |  | FOXA1 |  |  | MYCN |  |  |  |
|  |  |  |  |  |  |  |  | TPD52 |  |  |  |  |  |  |  |  |  |  |  |  |  |  |  |  |  | RAD51 |  |  |  |
|  |  |  |  |  |  |  |  | EGFR |  |  |  |  |  |  |  |  |  |  |  |  |  |  |  |  |  | PDL1 |  |  |  |
|  |  |  |  |  |  |  |  |  |  |  |  |  |  |  |  |  |  |  |  |  |  |  |  |  |  | AXL |  |  |  |
|  |  |  |  |  |  |  |  |  |  |  |  |  |  |  |  |  |  |  |  |  |  |  |  |  |  | CCNE1 |  |  |  |
|  |  |  |  |  |  |  |  |  |  |  |  |  |  |  |  |  |  |  |  |  |  |  |  |  |  | PEBP4 |  |  |  |
|  |  |  |  |  |  |  |  |  |  |  |  |  |  |  |  |  |  |  |  |  |  |  |  |  |  | MET |  |  |  |
|  |  |  |  |  |  |  |  |  |  |  |  |  |  |  |  |  |  |  |  |  |  |  |  |  |  | PDGFRA |  |  |  |
|  |  |  |  |  |  |  |  |  |  |  |  |  |  |  |  |  |  |  |  |  |  |  |  |  |  | PDGFRB |  |  |  |
|  |  |  |  |  |  |  |  |  |  |  |  |  |  |  |  |  |  |  |  |  |  |  |  |  |  | LyGDI |  |  |  |
| **Breast cancer** | | | | | | | | | | | | | | | | | | | | | | | | | | | | | |
| HN1 | HMGA2 | Notch1 | GIT1 | WT1 | CRKL | ROCK1 | KRAS | LMNB1 |  | Msi1 | MAPK7 | IRS1 | PIM1 | IGF1R | IRS1 | BIRC6 | MTDH | ELAVL1 | LIMK1 | RhoA | CDK8 | JAG1 | FOXA1 | Prrx2 | MKL1 | SIRT1 | EZH2 | CCND1 | Lin28 |
| FOXA1 | SALL4 | TOP2a |  |  | Foxf2 | PIK3CA |  | LMNA |  |  | LIMK1 | FASN |  | HOXA1 |  |  | Notch1 |  | FUT4 |  | PTGS2 | CCND1 | Six1 |  |  | BCL2 | ROCK1 |  | HMGA1 |
|  | Twist1 |  |  |  | KRAS |  |  |  |  |  | ERK5 | HMGCR |  | mTOR |  |  | Eya2 |  | PRKCA |  | SLC7A11 |  | JAK2 |  |  | RTCB | MYO10 |  | CCR7 |
|  |  |  |  |  | HMGB1 |  |  |  |  |  | CXCR4 | ACACA |  |  |  |  | Vim |  |  |  | TNKS1BP1 |  |  |  |  | TWIST1 | Met |  | PARP1 |
|  |  |  |  |  |  |  |  |  |  |  | MMP9 | CYP27B1 |  |  |  |  |  |  |  |  | CPSF7 |  |  |  |  | NOTCH1 |  |  |  |
|  |  |  |  |  |  |  |  |  |  |  | Kras | CCNE1 |  |  |  |  |  |  |  |  | COL12A1 |  |  |  |  | ZEB1 |  |  |  |
|  |  |  |  |  |  |  |  |  |  |  | VIM | Ccnd1 |  |  |  |  |  |  |  |  |  |  |  |  |  | GFRA3 |  |  |  |
|  |  |  |  |  |  |  |  |  |  |  | ERBB3 | RAF1 |  |  |  |  |  |  |  |  |  |  |  |  |  | SEMA4B |  |  |  |
|  |  |  |  |  |  |  |  |  |  |  | DNMT3A |  |  |  |  |  |  |  |  |  |  |  |  |  |  | MYB |  |  |  |
|  |  |  |  |  |  |  |  |  |  |  |  |  |  |  |  |  |  |  |  |  |  |  |  |  |  | CREB1 |  |  |  |
|  |  |  |  |  |  |  |  |  |  |  |  |  |  |  |  |  |  |  |  |  |  |  |  |  |  | MYC |  |  |  |
|  |  |  |  |  |  |  |  |  |  |  |  |  |  |  |  |  |  |  |  |  |  |  |  |  |  | LEF1 |  |  |  |
|  |  |  |  |  |  |  |  |  |  |  |  |  |  |  |  |  |  |  |  |  |  |  |  |  |  | FAM76A |  |  |  |
|  |  |  |  |  |  |  |  |  |  |  |  |  |  |  |  |  |  |  |  |  |  |  |  |  |  | CARKL |  |  |  |
|  |  |  |  |  |  |  |  |  |  |  |  |  |  |  |  |  |  |  |  |  |  |  |  |  |  | REM2 |  |  |  |
|  |  |  |  |  |  |  |  |  |  |  |  |  |  |  |  |  |  |  |  |  |  |  |  |  |  | ErbB2 |  |  |  |
|  |  |  |  |  |  |  |  |  |  |  |  |  |  |  |  |  |  |  |  |  |  |  |  |  |  | PRKD1 |  |  |  |
|  |  |  |  |  |  |  |  |  |  |  |  |  |  |  |  |  |  |  |  |  |  |  |  |  |  | LDHA |  |  |  |
|  |  |  |  |  |  |  |  |  |  |  |  |  |  |  |  |  |  |  |  |  |  |  |  |  |  | Wnt1 |  |  |  |
|  |  |  |  |  |  |  |  |  |  |  |  |  |  |  |  |  |  |  |  |  |  |  |  |  |  | TPD52 |  |  |  |
|  |  |  |  |  |  |  |  |  |  |  |  |  |  |  |  |  |  |  |  |  |  |  |  |  |  | LMTK3 |  |  |  |
|  |  |  |  |  |  |  |  |  |  |  |  |  |  |  |  |  |  |  |  |  |  |  |  |  |  | SRC |  |  |  |
|  |  |  |  |  |  |  |  |  |  |  |  |  |  |  |  |  |  |  |  |  |  |  |  |  |  | CDK4 |  |  |  |
|  |  |  |  |  |  |  |  |  |  |  |  |  |  |  |  |  |  |  |  |  |  |  |  |  |  | HDAC1 |  |  |  |
|  |  |  |  |  |  |  |  |  |  |  |  |  |  |  |  |  |  |  |  |  |  |  |  |  |  | HDAC7 |  |  |  |
|  |  |  |  |  |  |  |  |  |  |  |  |  |  |  |  |  |  |  |  |  |  |  |  |  |  | FOSL1 |  |  |  |
|  |  |  |  |  |  |  |  |  |  |  |  |  |  |  |  |  |  |  |  |  |  |  |  |  |  | Msi1 |  |  |  |
|  |  |  |  |  |  |  |  |  |  |  |  |  |  |  |  |  |  |  |  |  |  |  |  |  |  | AXL |  |  |  |
